# Arthropods and the evolution of RNA viruses

**DOI:** 10.1101/2021.05.30.446314

**Authors:** Tianyi Chang, Junya Hirai, Brian P. V. Hunt, Curtis A. Suttle

## Abstract

Many viruses of arthropods also infect other organisms including humans, sometimes with devastating consequences. Yet, for the vast diversity of arthropods, their associated viruses remain unexplored. Here, we mined meta-transcriptomes from 711 arthropod species, including insects, arachnids, myriapods, and crustaceans, and uncovered more than 1400 previously unknown RNA viruses, representing 822 novel evolutionary groups at a level between species and genus. These newly found viral groups fill major evolutionary gaps within the five branches of RNA viruses, bridging the evolution of viruses infecting early and later diverging eukaryotes. Additionally, co-phylogenetic analysis implies that RNA viruses of arthropods commonly co-evolved with their hosts. Our analyses indicate that arthropods have played a central role in the macroevolution of RNA viruses by serving as reservoirs in which viruses co-evolved with arthropods while being exchanged with a vast diversity of organisms.

---

Arthropods first appeared over 500 million years ago ^1,2^ and were among the first animals that pioneered terrestrial and freshwater ecosystems ^3^, during which time they formed close ecological relationships with plants and animals ^4^. In terrestrial ecosystems alone, there are estimated to be about 6.8 million arthropod species, far exceeding the sum of predicted species in the other three kingdoms comprising eukaryotes ^5,6^. Associated with these arthropods is an enormous diversity of viruses, some of which cause serious diseases in humans, livestock, and crops, at times with devastating effects ^7,8^. The viruses that are shared between arthropods and their plant and vertebrate hosts have gone through a co-evolutionary dance in which viruses that originally only infected arthropods acquired traits that also allowed them to replicate in plants ^9^ and vertebrates ^10^. Through this process, arthropods facilitated a massive expansion of the genetic diversity of RNA viruses that infect plants and animals.

## Results

### Revealing previously unknown diversity of RNA viruses associated with arthropods

Here, we explore the genetic diversity of RNA viruses associated with arthropods to determine the evolutionary relationships between these viruses and those that infect plants, fungi, and vertebrates. We did this by collecting representative gene sequences of the RNA-dependent RNA polymerase (RdRp), the hallmark gene for all RNA viruses, except retroviruses, for representatives of all the established families of RNA viruses (ICTV, 2019). We used this database to search for similar sequences in the Arthropoda subset of the Transcriptome Shotgun Assembly (TSA) database at NCBI. In addition, we performed meta-transcriptomic sequencing on marine copepods from five genera. In total, 1833 RNA viral genomes were retrieved from 29 orders of Hexapoda, 13 orders of Crustacea, and six orders of Chelicerata. Of these, more than 76 % belong to undescribed viruses, encompassing 822 previously unknown evolutionary groups (75% amino acid identity).

To differentiate true viruses from endogenous virus elements (EVEs), we applied a filtering approach based on genomic features of EVEs ^11,12^. Briefly, contigs were excluded if they carried interrupted viral genes or contained genes that were closely related to those of eukaryotes, EVEs, or transposable elements. The remaining virus-like sequences were compared with the genome sequences of their associated hosts and removed if they matched perfectly. However, only about 30% (519) of the newly found arthropod-associated RNA viruses (AARVs) had host genomes available; within these genomes three EVE-like contigs were identified, corresponding to 0.58% of the putative viruses that we identified in the meta-transcriptomes of these arthropods. Assuming these results are representative of those of other arthropods, only a fraction of a percent of the putative viruses that we identified were likely to be EVEs.

To resolve the evolutionary relationships among novel AARVs and established viral groups, we used inferred amino-acid sequences of the RdRp genes to construct maximum-likelihood phylogenetic trees of the 19 taxonomic groups of viruses in which AARVs are prominent (Fig. 1). This analysis shows that AARVs have been deeply involved in the evolution of major groups of RNA viruses, ranging from the presumptive earliest (*Lenarviricota*) to the latest diverging groups (*Negarnaviricota*). Overall, previously undescribed AARV lineages fill major gaps between established viral families and genera, and together with arthropod-borne plant and vertebrate viruses, make up a significant proportion of the diversity in major evolutionary groups of RNA viruses.

**Figure 1.**
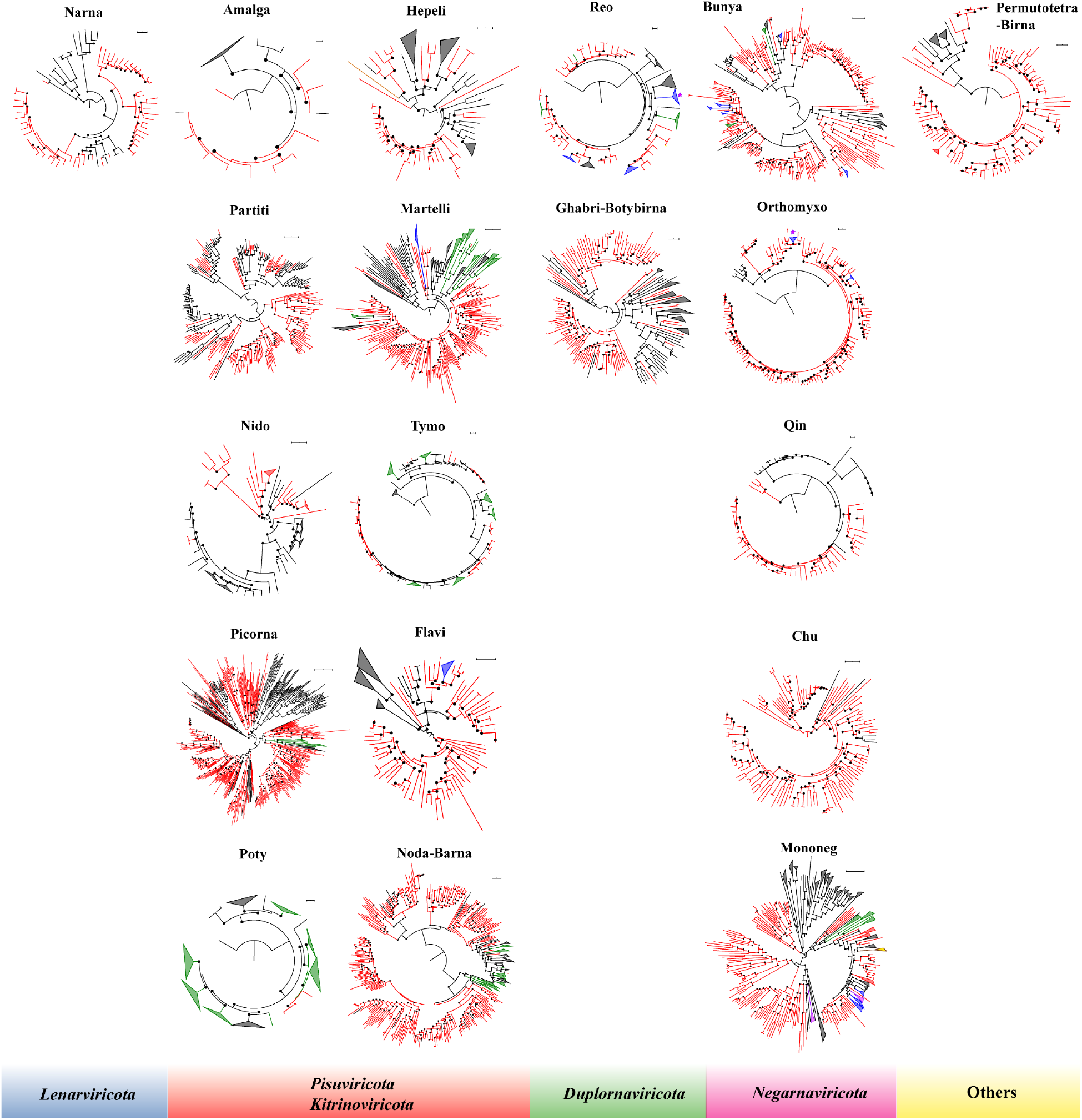
Phylogenetic analysis of 19 evolutionary groups in which AARVs reveals enormous diversity and shows the integral role of AARVs in RNA virus evolution. In each clade, the dataset comprises AARVs reported in this study and viruses with similar RdRp domains, as well as representatives of established genera within each clade. Phylogenies are derived from inferred amino-acid sequences of full length RdRps using maximum-likelihood. The phylogenetic trees are categorized into major evolutionary groups according to the ‘megataxonomy’ of RNA viruses ^13^. The branches within each phylogenetic tree are colored based on host range as follows: red, viruses discovered in arthropods; green, arthropod-borne plant viruses; blue, arthropod-borne mammal viruses; pink, arthropod-borne bird viruses; yellow, viruses found in arthropods and reptiles; black, viruses not detected in arthropods. In addition, viruses in the families *Reoviridae* and *Orthomyxoviridae* that encompass both arthropod-borne mammal viruses and arthropod-borne bird viruses are marked with asterisks. Branches are collapsed into genera. SH-aLRT branch support greater than 0.7 is shown by solid circles. Each scale bar indicates 0.5 amino-acid substitutions per site. The phylogenetic trees are mid-point rooted. *Narnaviridae* (Narna); *Amalgaviridae* (Amalga); *Partitiviridae* (Partiti); *Nidovirales* (Nido); *Picornaviridae* (Picorna); *Potyviridae* (Poty); *Hepelivirales* (Hepeli); *Martellivirales* (Martelli); *Tymoviridae* (Tymo); *Flaviridae* (Flavi); *Nodaviridae, Luteoviridae, Tombusviridae, Sobemovirus*, and *Barnaviridae* (Noda-Barna); *Reovirales* (Reo); *Ghabrivirales* and *Botybirnavirus* (Ghabri-Botybirna); *Bunyavirales* (Bunya); *Orthomyxoviridae* (Orthomyxo); *Qinviridae* (Qin); *Chuviridae* (Chu); *Mononegvirales* (Mononeg); *Permutotetraviridae* and *Birnaviridae* (Permutotetra-Birna).

Our analyses revealed that AARVs occurred in 12 major evolutionary groups of viruses. Viruses in the *Picornavirales* were most common and comprised the highest diversity of AARVs, and included 332 viruses from 14 orders of insects, nine orders of crustaceans, and five orders of arachnids (Fig. 2 and Suppl. Fig. 1). However, there were also 11 other major evolutionary groups of AARVs outside the *Picornavirales* in which between 57 and 299 viruses were identified (Suppl. Fig. 1); these encompassed viruses in the Noda-Barna (299), *Bunyavirales* (218), *Martellivirales* (156), *Mononegvirales* (146), *Partitiviridae* (133), *Orthomyxoviridae* (109), *Sobelivirales* (108), *Ghabrivirales* (89), *Chuviridae* (67), Permutotetra-Birna (60), and *Flaviviridae* (57). These groups likely encompass most of the RNA virus diversity in arthropods, as the results are based on searches for RdRp sequences of all taxonomic groups of RNA viruses in representative meta-transcriptomes from all arthropod subphyla.

**Figure 2.**
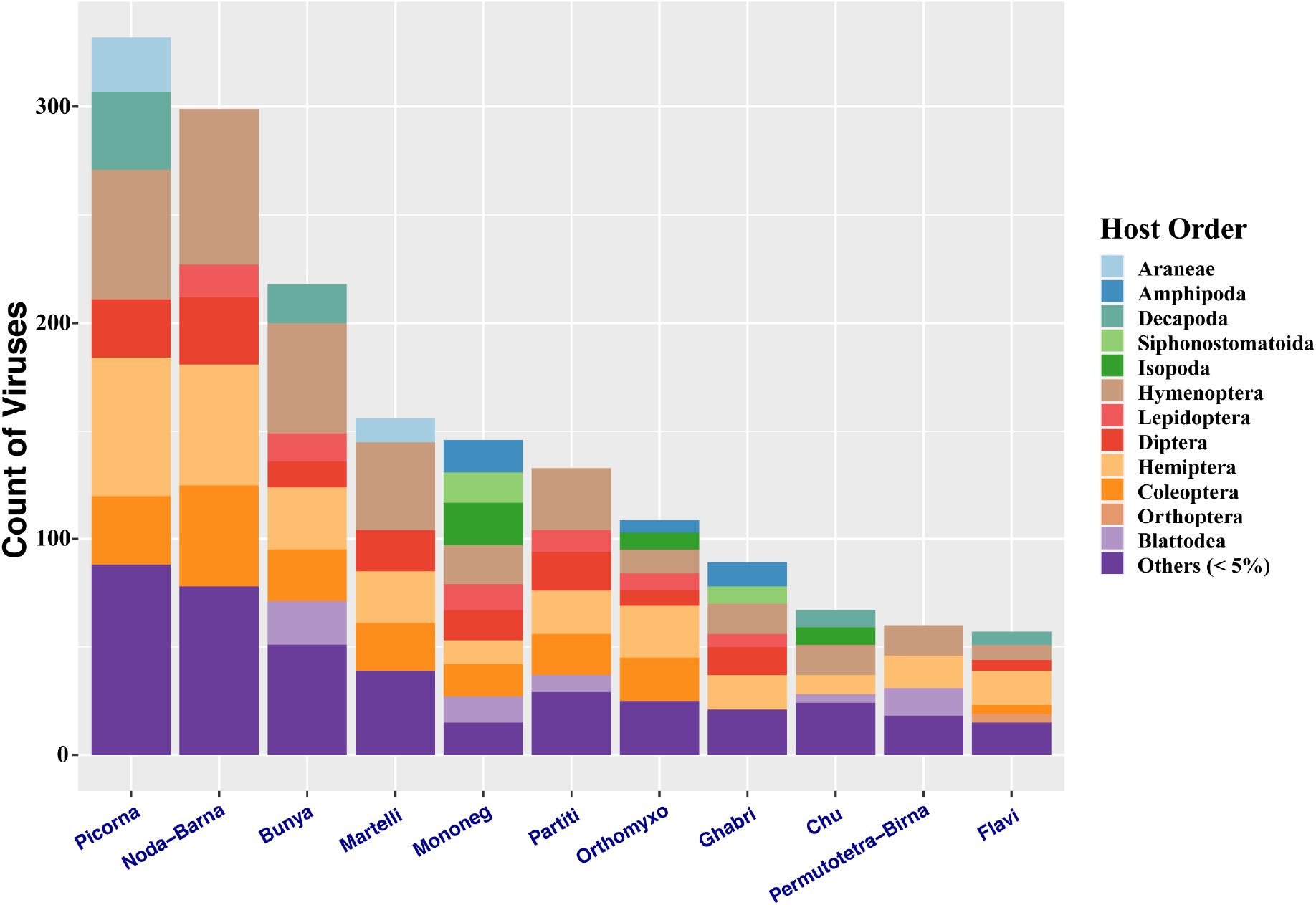
Major groups of RNA viruses associated with arthropods uncovered in the TSA database and in marine copepods. The number of RNA viruses binned by taxonomic group is shown for the different orders of arthropods with which they are associated. Orders of arthropods in which the RNA viruses were found are color coded, with orders that were associated with less than 5% of the relative abundance of viruses being assigned to Others. *Picornaviridae* (Picorna); *Nodaviridae, Luteoviridae, Tombusviridae, Sobemovirus*, and *Barnaviridae* (Noda-Barna); *Bunyavirales* (Bunya); *Martellivirales* (Martelli); *Mononegvirales* (Mononeg); *Partitiviridae* (Partiti); *Orthomyxoviridae* (Orthomyxo); *Ghabrivirales* (Ghabri); *Chuviridae* (Chu); *Permutotetraviridae* and *Birnaviridae* (Permutotetra-Birna); *Flaviviridae* (Flavi).

Our analysis significantly expands the known diversity of AARVs within major groups of RNA viruses as well as arthropod taxa with which the viruses are associated. The newly discovered viruses account for 18% to 55% of the total AARVs (75% amino-acid identity) for the *Nidovirales* (55%), Permutotetra-Birna (52%), *Qinviridae* (43%), *Orthomyxoviridae* (38%), *Hepelivirales* (30%), *Bunyavirales* (29%), *Chuviridae* (29%), *Ghabrivirales* (29%), Noda-Barna (28%), *Tymoviridae* (28%), *Mononegvirales* (24%), *Partitiviridae* (23%), *Reoviridae* (23%), *Flaviviridae* (23%), *Martellivirales* (22%), *Narnaviridae* (22%), and *Picornavirales* (18%) (Suppl. Fig. 2a). Likewise, a remarkable number of associated hosts of AARVs was revealed, accounting for 25% (*Reoviridae*) to 71% (Permutotetra-Birna) of hosts at the genus level (Suppl. Fig. 2b).

### Co-evolution between viruses and arthropods fueled the macroevolution of RNA viruses

The phylogenetic data highlight that the evolution of AARVs in arthropods is typically monophyletic within major evolutionary groups, such as families and orders (Fig. 1), and distinct from the evolution of viruses that infect other organisms including protists, fungi, and vertebrates. The pattern of monophyletic groups of viruses within related groups of arthropods (Suppl. Figs. 3-21) is consistent with virus-host co-evolution. Another indication of virus-host co-evolution is the similarity of the virus sequences to EVEs in arthropod genomes that are genetic remnants of ancient viral infections. Indeed, most AARVs share significant sequence homologies with EVEs in arthropods (Suppl. Figs. 3-21), implying deep evolutionary relationships between arthropods and major groups of RNA viruses, and that arthropods are the natural hosts of these viruses.

We further tested the congruency between the phylogenies of RNA viruses and the arthropods they infect. To do this, viral lineages related to the families *Narnaviridae, Flaviviridae, Partitiviridae, IFlaviviridae, Dicistroviridae, Orthomyxoviridae*, and *Chuviridae* were selected. These taxonomic groups cover the five major evolutionary branches of RNA viruses ^13^. Random permutation tests showed that the phylogenetic tree of arthropod viruses in these lineages is congruent with that of their hosts (p < 0.05) (Supplementary Tables 1 and 2), reflecting the co-evolution of RNA viruses with their arthropod hosts.

### Arthropods facilitated the diversification of RNA viruses through horizontal virus transmission

In addition to virus-host co-evolution, there is strong evidence for horizontal transmission of some AARVs between distantly related arthropods. For example, some negative-sense and positive-sense single-stranded RNA viruses infect both honeybees and their parasitic mites ^14–16^, while viruses related to the *Tymoviridae, Martellivirales, Picornavirales, Partitiviridae*, and *Orthomyxoviridae* were transmitted between spiders and insects (Suppl. Figs. 4, 6, 12, 16, and 21). As well, we found closely related partitiviruses in honeybees and mites (Suppl. Fig. 16), and dicistroviruses in a bird mite (*Dermanyssus gallinae*) and bird louse (*Menopon gallinae*) (Suppl. Fig. 12).

Additionally, our analysis supports that some RNA viruses that infect fungi, plants, or vertebrates originated from those of arthropods. For example, plant viruses assigned to the families *Tymoviridae, Kitaviridae, Reoviridae* (*Oryzavirus* and *Fijivirus*), *Tospoviridae*, and *Phenuiviridae* (*Tenuivirus*) (Suppl. Figs. 4, 6, 15, and 20) are all nested within clades of AARVs, consistent with these plant viruses being horizontally transmitted from arthropods through ecological interactions. It is unlikely that the AARVs are plant-specific; rather, they are most likely harboured by arthropods, since they are closely related to arthropod EVEs, and are associated with a wide spectrum of arthropods, including crustaceans that are unlikely to interact with plants. Likewise, fungal viruses in the *Reoviridae* (*Mycoreovirus*) (Suppl. Fig. 15), and vertebrate viruses in the *Togaviridae, Flaviviridae* (*Flavivirus*), *Reoviridae* (*Seadornavirus*), *Rhabdoviridae, Phenuiviridae* (*Phlebovirus, Bandavirus*, and *Phasivirus*), *Nairoviridae* (*Orthonairovirus*), *Peribunyaviridae* (*Orthobunyavirus*), and *Orthomyxoviridae* (*Thogotovirus* and *Quaranjaviru*s) (Suppl. Figs. 6, 7, 15, 19, 20, and 21) are all embedded within clades of arthropod viruses, indicating these viruses were also derived from arthropods.

### AARVs are linked to the evolution of genome organization and size in RNA viruses

We found that AARVs have had a major role in the genome diversification of RNA viruses. Members of the *Picornavirales* are of particular interest in that they encompass the highest diversity of AARVs, infect a broad range of eukaryotes, and are highly variable in genome size and structure. Indeed, we identified 21 different genomic architectures of arthropod-associated picornavirus-like viruses, representing a wide phylogenetic distribution (Fig. 3). Although generally comprising the same set of genes, the genomes of arthropod-associated picornavirus-like viruses underwent frequent rearrangements, sometimes losing or gaining structural genes, as well as changing the number of open reading frames and genome size. Non-parametric one-way ANOVA showed that genome size differs significantly among evolutionary groups (p < 1e-15). However, within each evolutionary group, the genome architecture is conserved in either one or two major forms, and the genome sizes are similar between architecture groups within the “Aquatic Picorna-like”, “Dicistroviridae-related”, and “Kelp fly virus related” (Mann-Whitney test, p > 0.5) (Supplementary Table 3).

**Figure 3.**
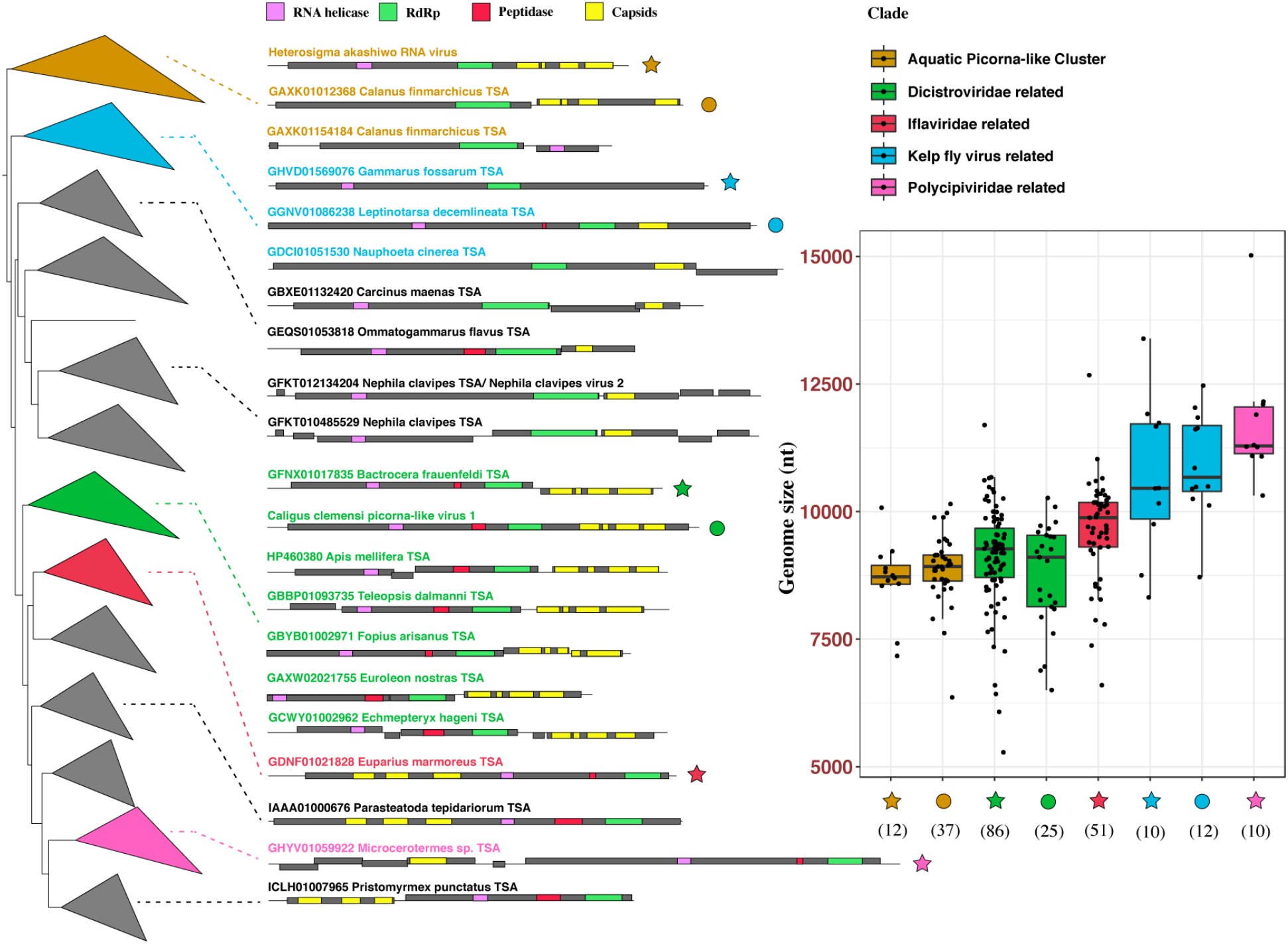
AARVs within the *Picornavirales* show consistent architecture and genome size within evolutionary groups. Phylogenetic tree inferred from RdRp domains of picornavirus-like viruses; branches are collapsed within each established virus clade. Genomic architectures of viruses are displayed for clades encompassing AARVs. In each clade, genomes with similar architecture are marked with either a star or circle, and the genome size distributions are shown in the boxplot. For each group in the boxplot, the interquartile range, median value, and non-outlier range are indicated by the box, solid line in the box, and whiskers, respectively; all the data is shown as individual points and the number of genomes for each architectural group is shown in the parenthesis. The major genomic architectures, as well as their corresponding clades and box plots are color coded. For genomes with multiple open reading frames (ORFs), the reading frames of the corresponding ORFs are indicated as follows: frame 1 (if the box representing the ORF is placed on the line representing the viral genome), frame 2 (if the box is placed under the line representing the viral genome), and frame 3 (if the box is placed above the line representing the viral genome). Functional domains within each ORF are color coded and indicated by the associated figure legends.

## Discussion

Our analyses demonstrate the central role of arthropods in the macroevolution of RNA viruses (Fig. 4). By comparing sequences of arthropod EVEs and AARVs, we show that arthropods have been both widely and deeply involved in RNA virus evolution. Furthermore, co-phylogenetic analysis between RNA viruses of arthropods and their hosts are consistent with virus and host co-evolution, implying that the enormous diversity of AARVs was a consequence of arthropod radiations. As arthropods built ecological connections in aquatic and terrestrial systems, they exchanged their viruses with a wide array of plants, fungi, and other animals, and facilitated the divergence of RNA viruses.

**Figure 4.**
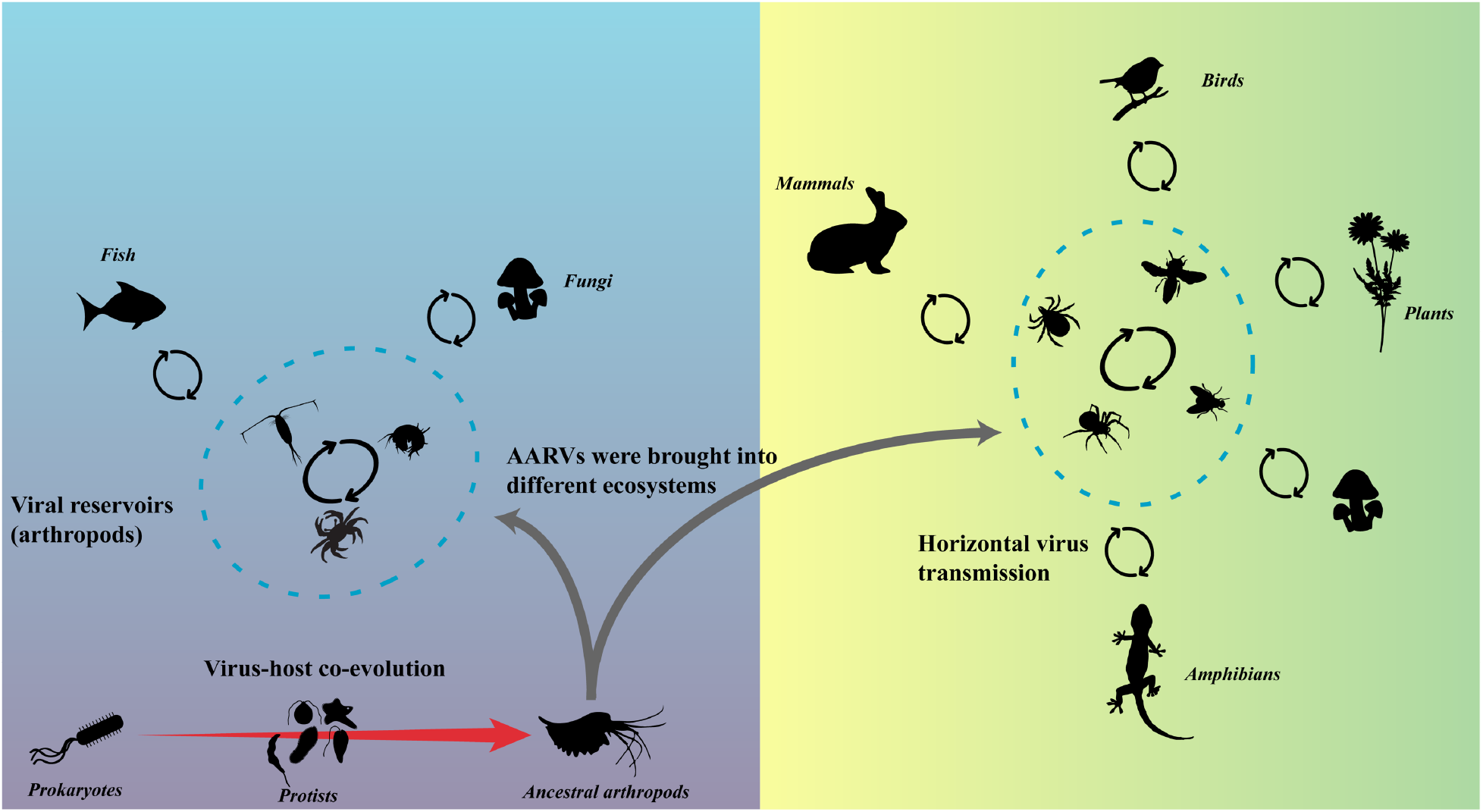
RNA virus macroevolution was shaped by arthropods. The red arrows represent long-term virus-host co-evolution, including RNA virus transmissions from bacteria to early eukaryotes, and from ancient to modern arthropods. The figure highlights the role of arthropods in connecting the evolution of RNA viruses between early and late diverging eukaryotes. The black circular arrows represent the transmission of viruses among arthropods and between arthropods and fungi, plants, and vertebrates through ecological connections in aquatic (left) and terrestrial (right) ecosystems.

It is well known that viruses can be exchanged between insects and plants, and between parasitic arthropods and vertebrates, including plant viruses vectored by thrips and whiteflies, and vertebrate viruses transmitted by mosquitos and ticks. Indeed, many of the plant and vertebrate RNA viruses that we identified that originated in arthropods have arthropod vectors, consistent with the horizontal acquisition of these viruses from arthropods. Likewise, in aquatic ecosystems, closely related viruses within the *Flaviviridae* and *Nodaviridae* occur in fish and crustaceans. For instance, closely related viruses were found in a crab and shark ^17^; as well, fish nodaviruses are within a group of viruses associated with a copepod, a barnacle, horseshoe crabs, and decapods (Suppl. Fig. 8). Additionally, viruses in the genus *Caligrhavirus* that infect sea lice (parasitic copepods of fish) are at the base of viruses in the genera *Perhabdovirus* and *Sprivivirus* (Suppl. Fig. 19), which infect fish. These phylogenetic relationships provide strong evidence of transmission of RNA viruses between fish and crustaceans.

There are also striking linkages between RNA viruses of arthropods and those of fungi for the five major branches of RNA viruses, implying that fungi and arthropods have intimate ecological relationships that have resulted in the exchange of viruses. For example, viruses in the *Mitoviridae* (*Lenarviricota*), *Gammapartitivirus* (*Pisuviricota*), *Alphaendornavirus* (*Kitrinoviricota*) and *Totiviridae* (*Duplornaviricota*) are all sisters to AARV clades (Suppl. Figs. 1, 6, 13, and 16). We also identified a novel rhabdo-like virus (*Negarnaviricota*) from a parasitic fungus of flies (*Entomophthora muscae*); the virus falls within a deep lineage of rhabdo-like viruses that likely infect arthropods (Suppl. Fig. 19). Additionally, our data show that some viruses of arthropods, plants, and phytopathogenic fungi were derived from the same ancestors in the *Tymovirales, Hepelivirales*, and *Partitiviridae* (Suppl. Figs. 4, 5, and 16), indicating that the evolution of these viruses likely resulted from ecological interactions among the three groups of organisms. Studies on three-way interactions among plants, fungi, and viruses have revealed different biological effects ^18,19^. Our findings suggest that ecological interactions among arthropods, plants, fungi, and viruses were important drivers of RNA virus evolution.

Several studies have opened our eyes to the massive diversity of AARVs ^20–24^, while others have pointed out the important role of arthropods in the evolution of RNA viruses ^9,20,25,26^. Our study emphasizes the intertwined evolutionary relationship between viruses within arthropods and other organisms intimately associated with them, and greatly expands the known diversity of AARVs, and the range of hosts with which they are associated. Additionally, we identified previously unknown lineages of viruses that likely infect arthropods of economic and ecological significance. These include viruses in sea lice (parasitic copepods) that have been implicated in the decline of wild salmon stocks and cause significant economic losses in fish aquaculture ^27,28^. As well, many previously unknown viruses were found to be associated with planktonic copepods, the most abundant arthropods on earth, and a critical link in marine foodwebs ^29^. We also discovered many viruses of ticks and mosquitos that are disease vectors of humans, livestock, and wild animals ^7^, and viruses of white flies, plant hoppers, and thrips, which transmit pathogens among many plants of economic and ecological importance ^8,30,31^. These findings highlight the massive diversity of viruses that are hidden in arthropods, and demonstrate the central role that arthropods have played in the evolution of viruses infecting multicellular life.

## Methods

### Virus discovery from the Transcriptome Shotgun Assembly (TSA) Database

To search for AARVs in the TSA database in an unbiased manner, we collected representative genomes from all the established viral families within the *Riboviria* except for members encoding reverse transcriptases (the *Revtraviricetes*). Predicted open reading frames from the representative viruses were predicted with ORFfinder (https://www.ncbi.nlm.nih.gov/orffinder/), and putative coding sequences for RdRps were identified by searching against the NCBI Conserved Domain Database ^32^. Since RdRp sequences can be highly divergent, they were translated from the representative viruses and used as probes to search against the Arthropoda subset of the TSA Database using TBLASTN. Contigs that shared significant sequence similarity with RdRp (*e*-value < 1e-5) were manually screened based on their completeness and authenticity; those shorter than 200 amino acids, as well as potential EVEs or chimeras, were excluded from further analysis.

### Virus discovery from marine copepods

We investigated the RNA virome in the following five genera of marine copepods: *Caligus clemensi, Cosmocalanus darwinii, Pleuromamma abdominalis, Euchaeta* spp., and *Eucalanus bungii*. Among them, *C. clemensi* is a parasite of fish and in the order Siphonostomatoida, while the other four species are pelagic copepods belonging to the order Calanoida.

Individuals of *C. clemensi* were collected from four species of salmon, coho (*Oncorhynchus kisutch*), chum (*O. keta*), pink (*O. gorbuscha*) and sockeye (*O. nerka*), as well as Pacific herring (*Clupea pallasii*). The fish were captured by purse seine at five sampling sites ranging from the southern Discovery Islands to northern Johnstone Strait, British Columbia, Canada, during early, mid, and late summer from 2015 to 2017 (Department of Fisheries and Oceans Canada permits: XR422015, XR922016, XR252017). The fish were sampled individually from the net, euthanized with Tricaine methanesulfonate (MS-222) ^33^ in accordance with UBC Animal Care Protocol A16-0101, following Canadian Council on Animal Care guidelines, and immediately frozen in liquid nitrogen. Motile-stage sea lice were picked off the fish and based on morphological features observed under a dissecting microscope identified by species, sex, and life stage, and immediately frozen at -80 °C until further processing.

Calanoid copepods were collected with a 100-µm or 250-µm mesh-size plankton net, and identified, selected, and then depurated for at least 3 h in 0.45-µm filtered seawater at ambient water temperature. After depuration, the copepods were immersed in RNA*later* (Ambion, Inc., Austin, TX) at 4 °C overnight, and then transferred to -80 °C until nucleic acids were extracted. Individuals of *E. bungii* were collected in the Strait of Georgia on 25 April 2018, while individuals of *P. abdominalis, Euchaeta* spp. and *C. darwinii* were collected in November 2019 from oligotrophic waters in the Southeast Pacific Ocean.

We used the kit Direct-zol RNA Miniprep Plus (Zymo Research Corporation) to perform total RNA extractions. For samples preserved in RNA*later*, excessive reagent was removed from copepods with Kimwipes™ in a sterile and RNase-free environment prior to extraction. Individuals (30 to 100) were pooled by species and homogenized in TRI reagent (Sigma-Aldrich) with a pellet pestle, and the total RNA extracted from the homogenate following the manufacturer instructions. The ribosomal RNA from eukaryotes and bacteria were removed using the Ribo-Zero Plus rRNA Depletion Kit (Illumina), and RNAseq libraries were constructed with either the NEB directional RNA preparation kit (*P. abdominalis, Euchaeta* spp., and *C. darwinii*) or the NEBNext Ultra RNA Library Prep Kit (*C. clemensi* and *E. bungii*). To ensure enough sequencing depth for virus discovery, only two to three libraries were pooled and sequenced per lane on the Illumina HiSeq platform (2 × 150 bp PE mode).

For each library, raw reads were trimmed, and quality checked with Trimmomatic v0.38 ^34^ and FastQC (http://www.bioinformatics.babraham.ac.uk/projects/fastqc/), respectively. Eukaryotic and bacterial rRNA were further removed from the post-QC reads using SortMeRNA v4.1 ^35^. We then assembled the remaining reads with both Trinity v2.8.4 ^36^ and rnaSPAdes v3.14.0 ^37^ and the resulting contigs were searched against the NCBI-nr database (downloaded in April 2019) using DIAMOND BLASTX v0.9.24.125 (*e* value cutoff: 1e-3) ^38^. To avoid false positives, only the sequences for which RdRp was the top hit were retained, and only those with a complete or near-complete coding sequence were kept for further analysis.

### Phylogenetic analysis and RdRp clustering

Phylogenies of RNA viruses were constructed from inferred amino-acid sequences for RdRp. Notably, for some viral groups, different fragments containing the RdRp gene were used to infer the phylogenetic trees among studies. To make our results more comparable with previous studies, we analyzed our RdRp sequences based on regions commonly used for phylogenetic analysis. Specifically, for viruses related to the *Reoviridae* and *Orthomyxoviridae*, only segments encoding RdRps were analyzed; for viruses belonged to the *Potyviridae*, the coding sequences of the entire RdRp containing polyprotein were used; and, for viruses in the *Picornavirales*, only sequences encoding the conserved domains of the RdRp were used for phylogenetic analysis. For reference RdRp sequences in each major evolutionary group, we included representatives from all the viral genera within each group, as well as other viruses that shared significant sequence similarity with the AARVs reported here.

To test the robustness of the methods used to construct the phylogenetic trees, we compared results obtained using different bioinformatic tools. These included T-Coffee, MUSCLE, and MAFFT for sequence alignment, different modes in TrimAl for alignment trimming, and RAXML-ng v9 and PhyML v3.0 for tree construction ^39–44^. In most cases the architectures of the phylogenetic trees were stable when generated using different approaches. However, for viruses in the *Picornavirales*, both the selection of the input RdRp sequences and the sequence alignment construction methods can affect the tree topology, likely due to the highly divergent RdRps sequences that occur within these viruses. Nevertheless, this does not affect our conclusions, since most AARVs fall within the same groups, and many of these viruses (e.g., “Kelp Fly Virus” related and “GFJG01075353 Eurypanopeus depressus TSA” related) consistently form deep-branching lineages that are at basal positions to other established viral groups.

To construct the final phylogenetic trees, we used CD-hit v4.8.1 ^45^ to dereplicate the input RdRp protein sequences, and then applied MUSCLE v3.8.31 and TrimAl v1.4 (strict mode) on the dereplicated sequences to construct the sequence alignment and quality screen, respectively. The trimmed alignments were manually checked in Jalview v2.11.1.4 ^46^ the phylogenetic trees were inferred using the Maximum-likelihood approach in PhyML v3.0. For each alignment, the best amino-acid substitution model was selected based on the Bayesian Information Criterion integrated in the Smart Model Selection algorithm ^47^. The final tree topology was determined by starting with 10 random trees and then searching with the Subtree-Pruning-and-Regraft method. Branch supports were measured using the Shimodaira–Hasegawa SH-aLRT algorithm, which is suggested to outperform standard bootstrap and is more robust to model violations ^48,49^.

We defined a virus as undescribed if its RdRp sequence did not show 100% identity to a known viral sequence. To identify novel evolutionary groups of RNA viruses, viral RdRp proteins were clustered with CD-hit v4.8.1 ^45^. The database of reference sequences encompassed representatives from all viral genera assigned to the *Riboviria* except the *Revtraviricetes*, as well as unclassified AARVs from recent studies ^22–24^. The inclusion of the newly discovered RdRp protein sequences to those of the database added 882 previously unknown groups of viruses (> 75% identity) at a level between species and genus. We took the same approach to estimate the number of newly identified groups of AARVs in each major groups of RNA viruses.

### Comparison of AARVs and EVEs of arthropods

To predict whether the newly discovered AARVs infect arthropods, and to differentiate viruses from EVEs, we downloaded all available arthropod genome assemblies from Whole Genome Shotgun projects (https://www.ncbi.nlm.nih.gov/Traces/wgs/) and used them as queries to perform DIAMOND BLASTX v0.9.24.125 searching for related AARV sequences. A genomic sequence was considered to be an EVE if it shared significant sequence similarity (*e*-value < 1e-5) with sequences of AARVs.

### Co-phylogenetic analysis between arthropods and viruses

We selected seven of the newly discovered AARV lineages to examine virus-host co-evolution. Viruses in these lineages likely infect arthropods since most share significant sequence similarity (*e*-value < 1e-30) with arthropod EVEs (DIAMOND BLASTX v0.9.24.125). The selected AARV lineages encompass or are sister groups to the *Narnaviridae* (n = 49), *IFlaviviridae* (n = 119), *Dicistroviridae* (n = 150), *Flaviviridae* (n = 86), *Partitiviridae* (n = 111), *Orthomyxoviridae* (n = 131), and *Chuviridae* (n = 119), covering Baltimore classes III, IV, and IV and the five major evolutionary branches of RNA viruses ^13^. The co-phylogenetic analyses were conducted by constructing matrices that consisted of host and virus evolutionary distances. For viruses, evolutionary distances were retrieved from the phylogenetic trees in R ^50^, and the evolutionary distances between different arthropod orders were obtained from previous studies based on their estimated time of divergence ^51–58^. We tested the congruence between phylogenies of viruses and their hosts based on the distance matrices using ParaFit ^59^ in the R package ape ^60^, and the p-value was calculated from 10,000 random permutations of each viral lineage.

### Construction and analysis of size variation of viral genomes

ORFs in viral genomes were predicted using ORFfinder (https://www.ncbi.nlm.nih.gov/orffinder/), and then annotated by BLAST searching in the NCBI Conserved Domains Database (https://www.ncbi.nlm.nih.gov/Structure/cdd/cdd.shtml). Since many novel viruses were found to contain highly divergent genes, the cutoff of *e*-value was set to 1, and only genes previously found in viruses were retained.

A linear model was used to test if there is a difference in genome size among evolutionary groups of picorna-like viruses. Then, a Shapiro-Wilk test was carried out to assess the normality of residuals of the regression as well as the genome sizes of viruses within each evolutionary group. Since the data were not normally distributed, we used Kruskal-Wallis and Mann-Whitney U tests to determine if genome size differs significantly among evolutionary and architectural groups of picorna-like viruses, respectively.

## Supporting information

Host-virus association of AARVs

## Data availability

Raw reads of marine copepods have been submitted to the Sequence Read Archive under accession number PRJNA700427.

The sequence alignments of viral RdRp proteins used to construct the phylogenetic trees are publicly available at https://dx.doi.org/10.6084/m9.figshare.14442806.

## Acknowledgements

We thank Amy Chan, Kevin Zhong, and all members of the Suttle lab, as well as Yanting Liu for their feedback throughout the project. We thank Gideon Mordecai for his review and feedback on the manuscript. We greatly appreciate the help of Julian Gan, Carly Janusson, and Brett Johnson as well as other members of the Hakai Institute for providing sea lice and zooplankton samples, and the crew of the Hakuho Maru for facilitating the collection of copepods from the southeast Pacific Ocean. This research was supported by a Discovery Grant from the National Science and Engineering Council of Canada (NSERC) to C.A.S., a scholarship from the Chinese Scholarship Council to T.C., and a fellowship and research grant to J.H. from the Japan Society for the Promotion of Science.

## Author contributions

T.C. and J.H. performed sample collection and pre-processing for pelagic copepods; B.P.V.H. organized and supervised the collection of sea lice; T.C. carried out data mining and analyses under the supervision of C.A.S; T.C. and C.A.S wrote the manuscript, which was edited by all authors.

