## Supplementary Figures for "Arthropods and the evolution of RNA viruses"

### Supplementary figures
# S1-S21

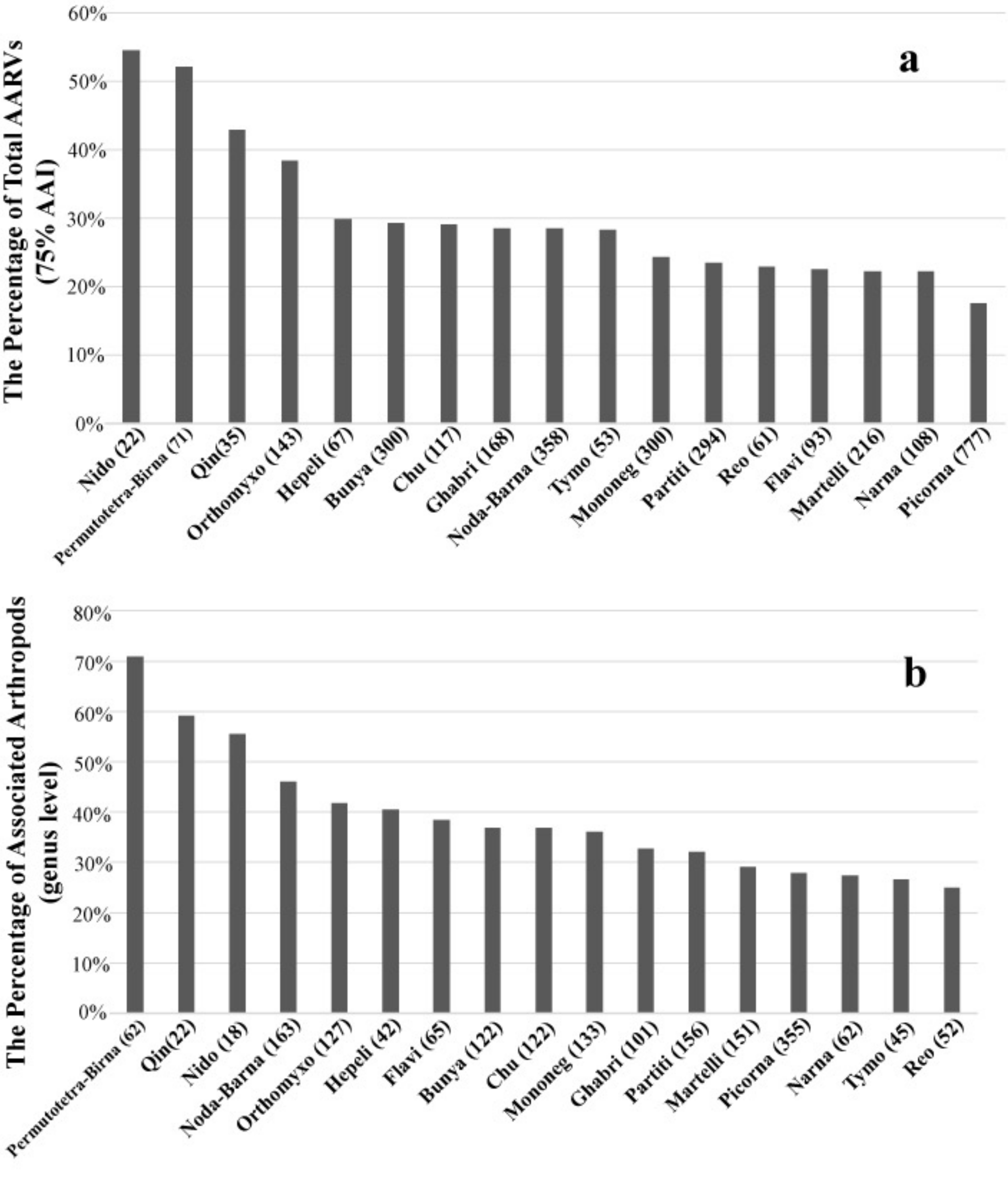


**Supplementary Figure 1. The percentages of AARVs identified in this study and newly identified genera of arthropods in which AARVs were found relative to the total known, binned by major taxonomic groups of viruses.**

Bars represent **(a)** the percentage of known AARVs contributed by this study clustered at 75% amino-acid identity for RdRp sequences, and **(b)** the newly identified arthropod genera in which the different types of viruses were found. The total number of non-redundant AARVs (< 75% identity) and genera of hosts are shown in parentheses for each viral group in **(a)** and **(b)**, respectively. *Picornaviridae* (Picorna); *Nodaviridae*, *Luteoviridae*, *Tombusviridae*, *Sobemovirus*, and *Barnaviridae* (Noda-Barna); *Bunyavirales* (Bunya); *Mononegvirales* (Mononeg); *Partitiviridae* (Partiti); *Martellivirales* (Martelli); *Ghabrivirales* (Ghabri); *Orthomyxoviridae* (Orthomyxo); *Chuviridae* (Chu); *Narnaviridae (*Narna); *Flaviviridae* (Flavi); *Permutotetraviridae* and *Birnaviridae* (Permutotetra-Birna); *Hepelivirales* (Hepeli); *Reovirales* (Reo); *Tymoviridae* (Tymo); *Qinviridae* (Qin); *Nidovirales* (Nido).

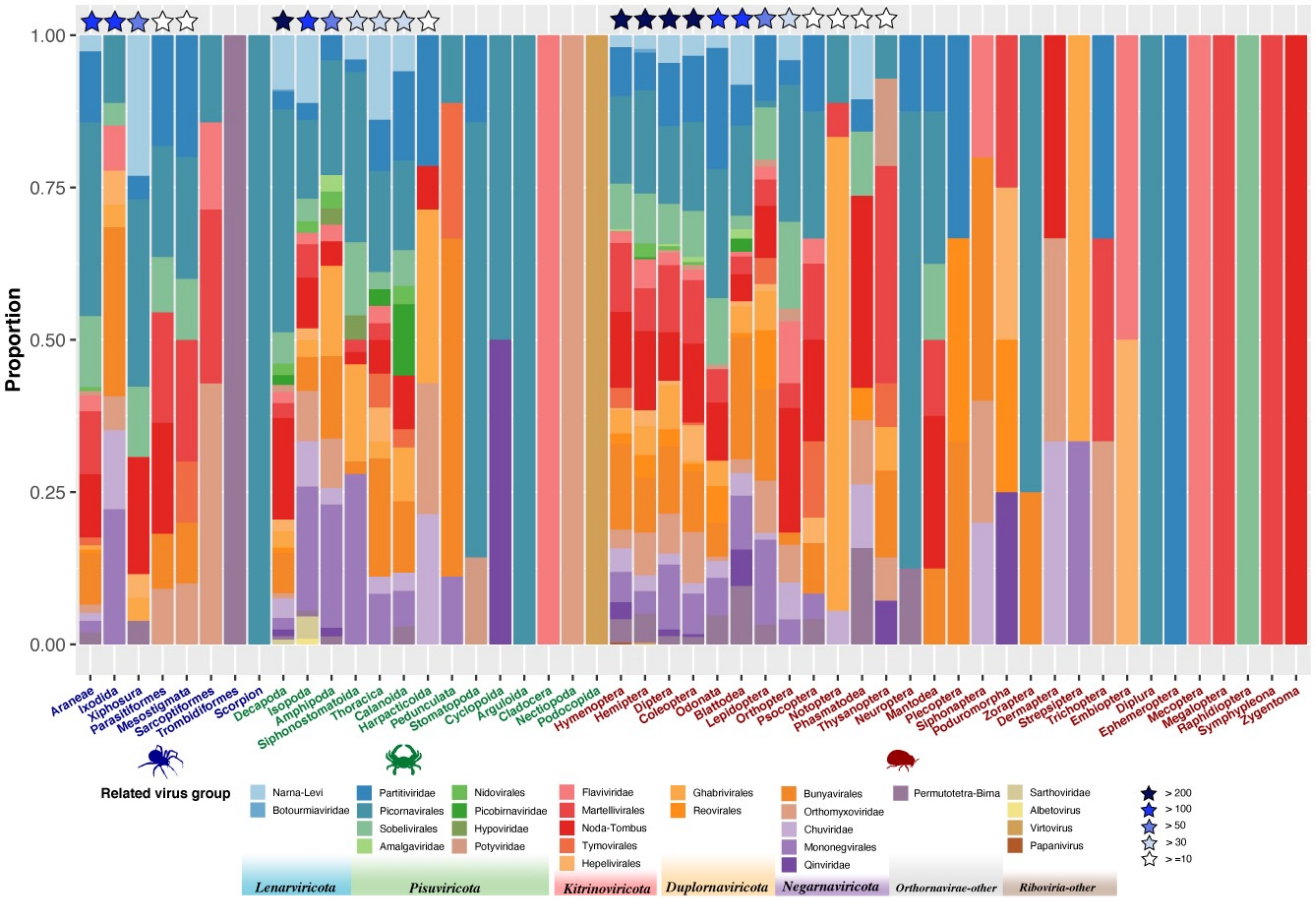


**Supplementary Figure 2: RNA virome structures in major arthropod orders.**

The figure shows the contribution of major RNA virus groups to virome of major arthropod order. Arthropod orders encompassing more than 10 RNA viruses are indicated by stars above the corresponding bars.

### (+) RNA viruses

- S3: Phylogeny of viruses related to the *Narnaviridae.*
- S4: Phylogeny of viruses related to the *Tymovirales*.
- S5: Phylogeny of viruses related to the *Hepelivirales*.
- S6: Phylogeny of viruses related to the *Martellivirales.*
- S7: Phylogeny of viruses related to the *Flaviviridae*.
- S8: Phylogeny of viruses related to the “Noda-Barna”.
- S9: Phylogeny of viruses related to the *Nidovirales*.
- S10: Phylogeny of viruses related to the *Potyviridae*.
- S11: Phylogeny of viruses related to the “Birna-Permutotetra”.
- S12: Phylogeny of viruses related to the *Picornavirales*.

#### Figure legends for all the phylogenies

Established viral groups (i.e. families and genera) are collapsed and color coded. Branches representing unclassified AARVs are colored in red (solid lines: novel AARVs found here; dotted lines: previously found AARVs), and orders of the associating arthropods are also color coded. SH-aLRT branch support greater than or equal to 0.9, less than 0.9 and greater than or equal to 0.8, less than 0.8 and greater or equal to 0.6 is shown by circles filled with black, grey, and white colors, respectively. AARVs which show significant sequence homologies to EVEs of arthropods are marked with red stars. The solid red stars represent AARVs with e values (BLASTn) lower than 1e-30, and the transparent red stars represent AAVs with e value fall between 1e-30 and 1e-5. If the majority of AARVs in a lineage shared significant sequence homologies with arthropod EVEs, the whole branch is marked with solid red stars and viruses which have no significant hits to the EVEs are indicated by empty stars on the corresponding branches. The scale bar indicates 0.2 amino acid substitutions per site. The phylogenetic tree is mid-point rooted.

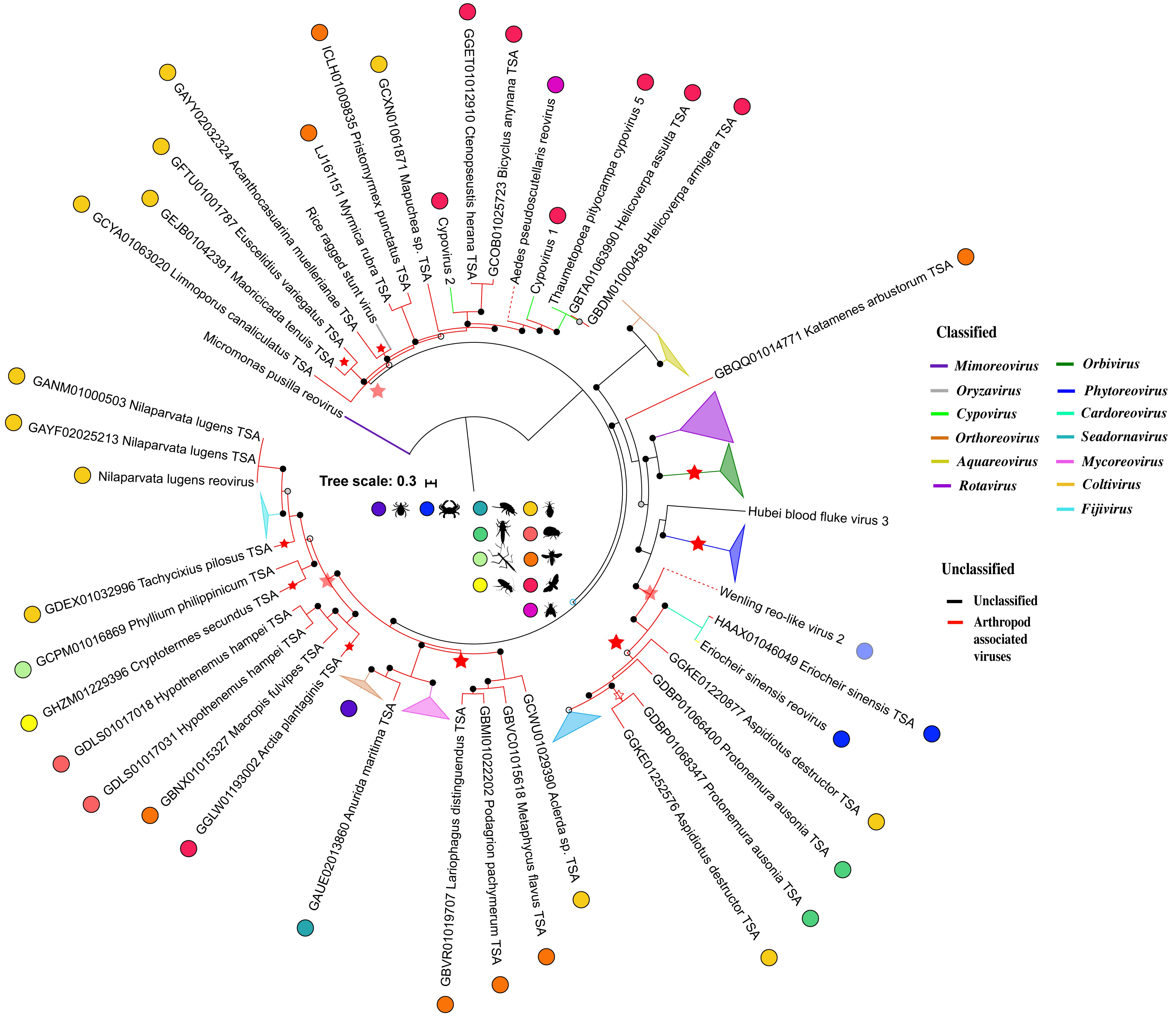


**Supplementary Figure 3: Phylogeny of viruses related to the *Narnaviridae.***

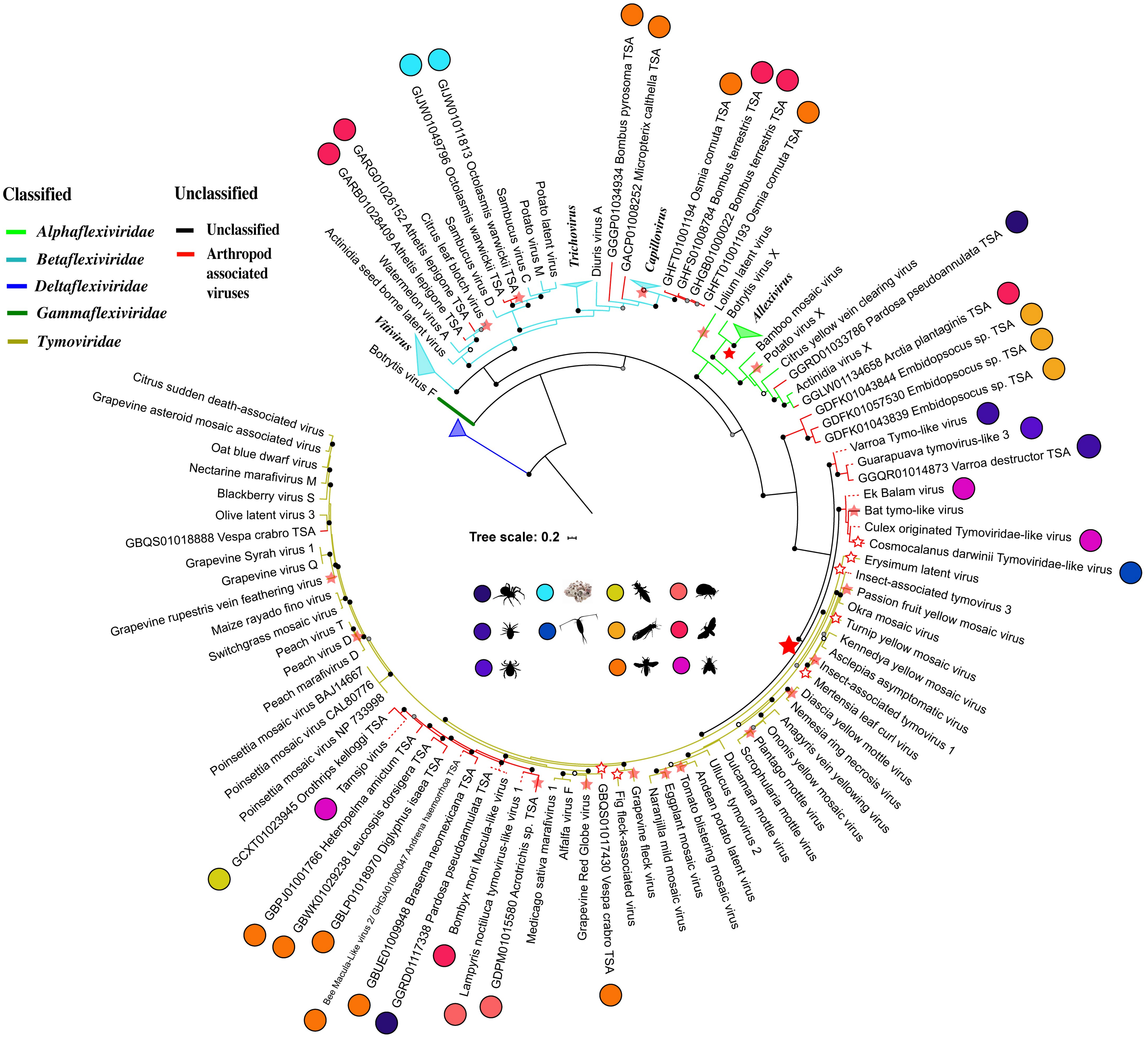


**Supplementary Figure 4: Phylogeny of viruses related to the *Tymovirales*.**

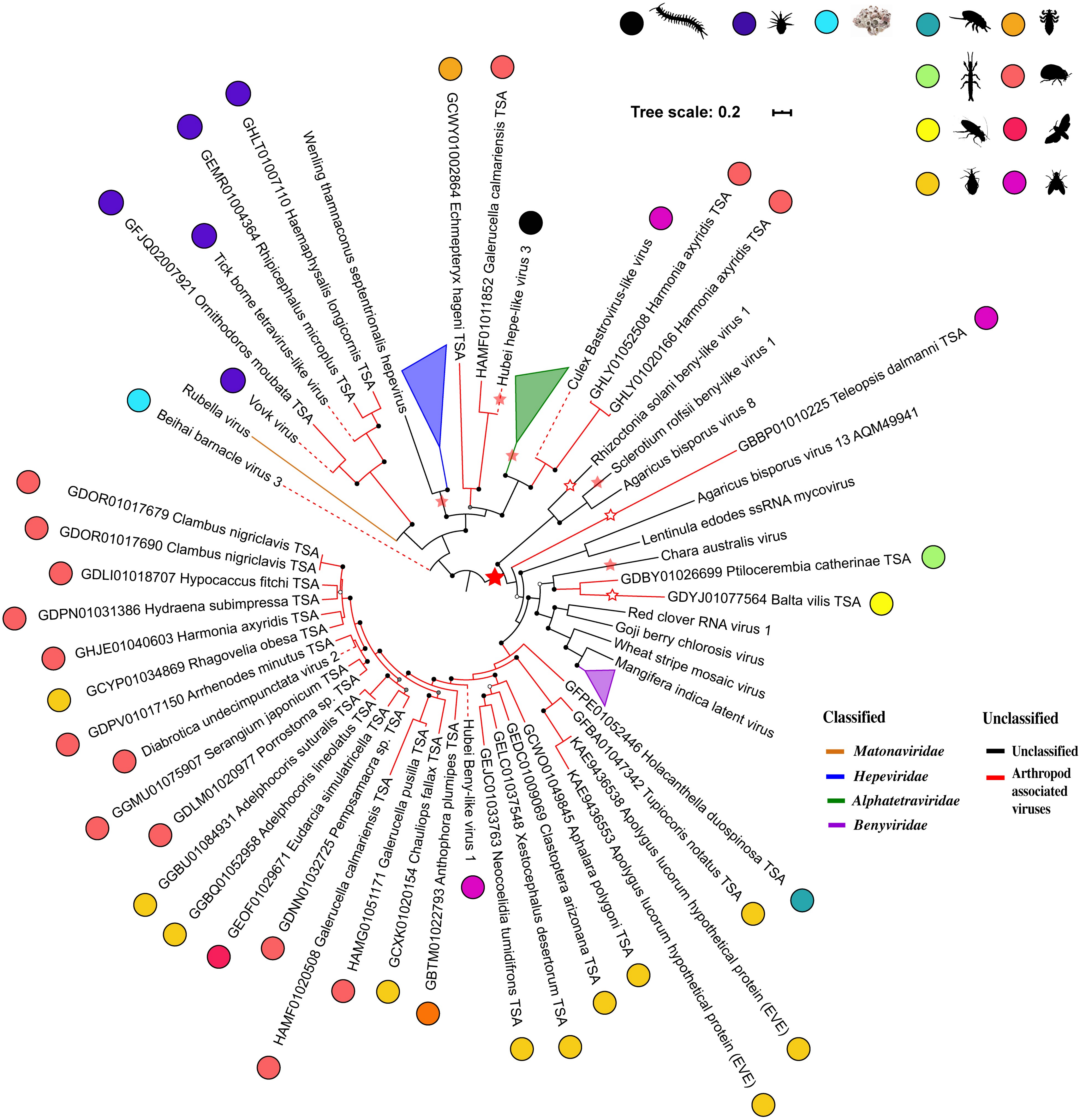


**Supplementary Figure 5: Phylogeny of viruses related to the *Hepelivirales*.**

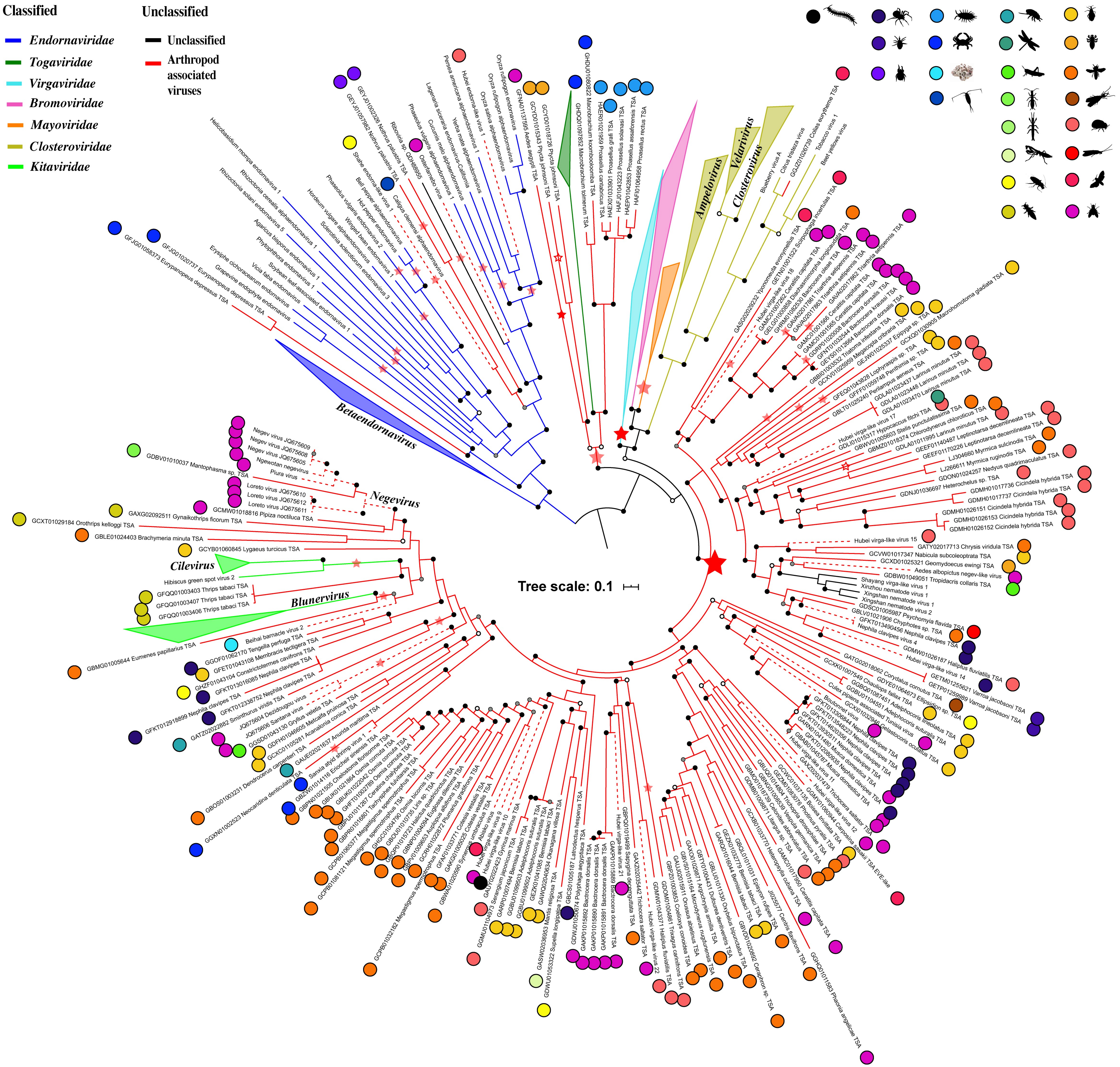


**Supplementary Figure 6: Phylogeny of viruses related to the *Martellivirales*..**

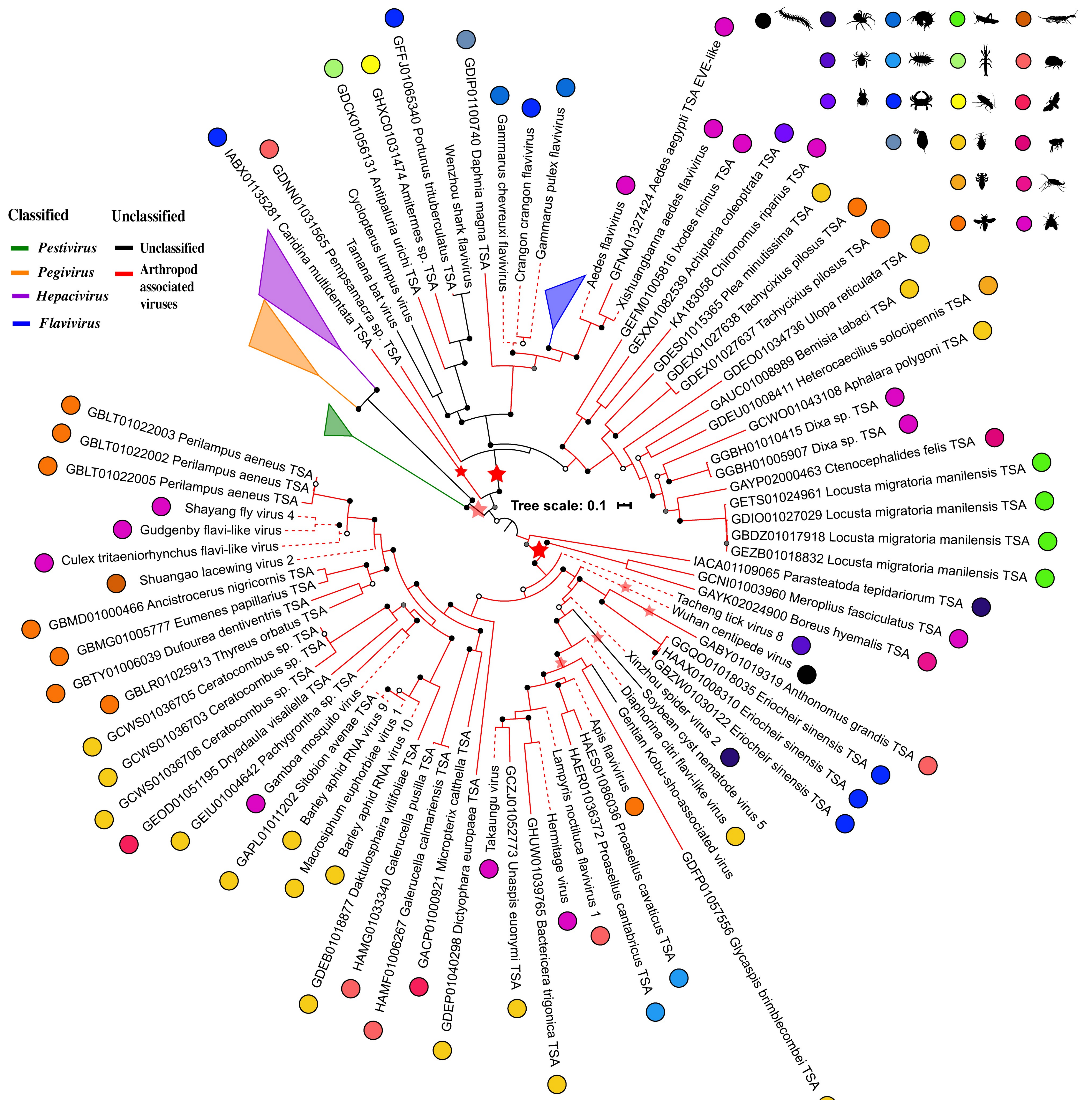


**Supplementary Figure 7: Phylogeny of viruses related to the *Flaviviridae*..**

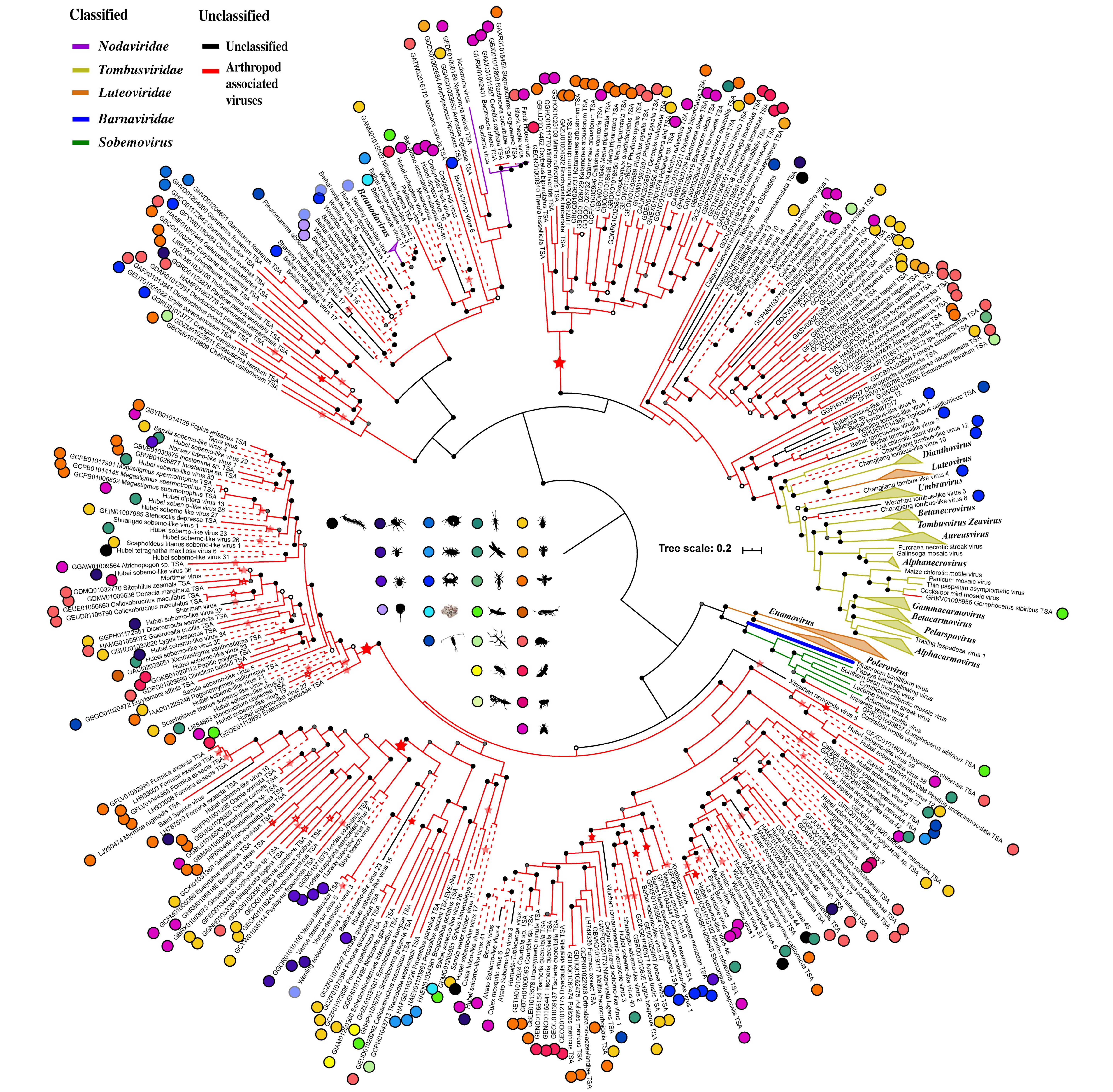


**Supplementary Figure 8: Phylogeny of viruses related to the "Noda-Barna".**

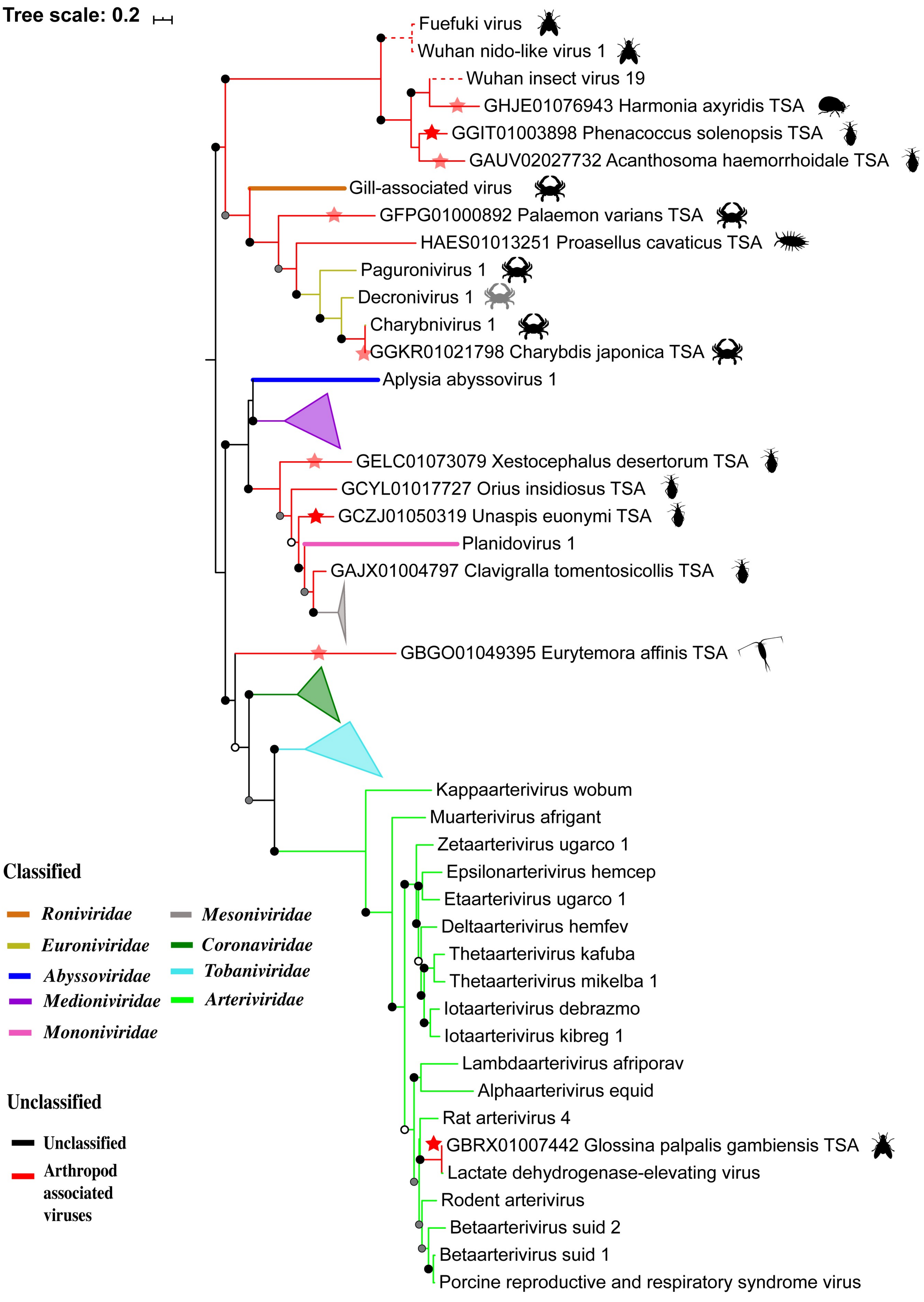


**Supplementary Figure 9: Phylogeny of viruses related to the *Nidovirales*..**

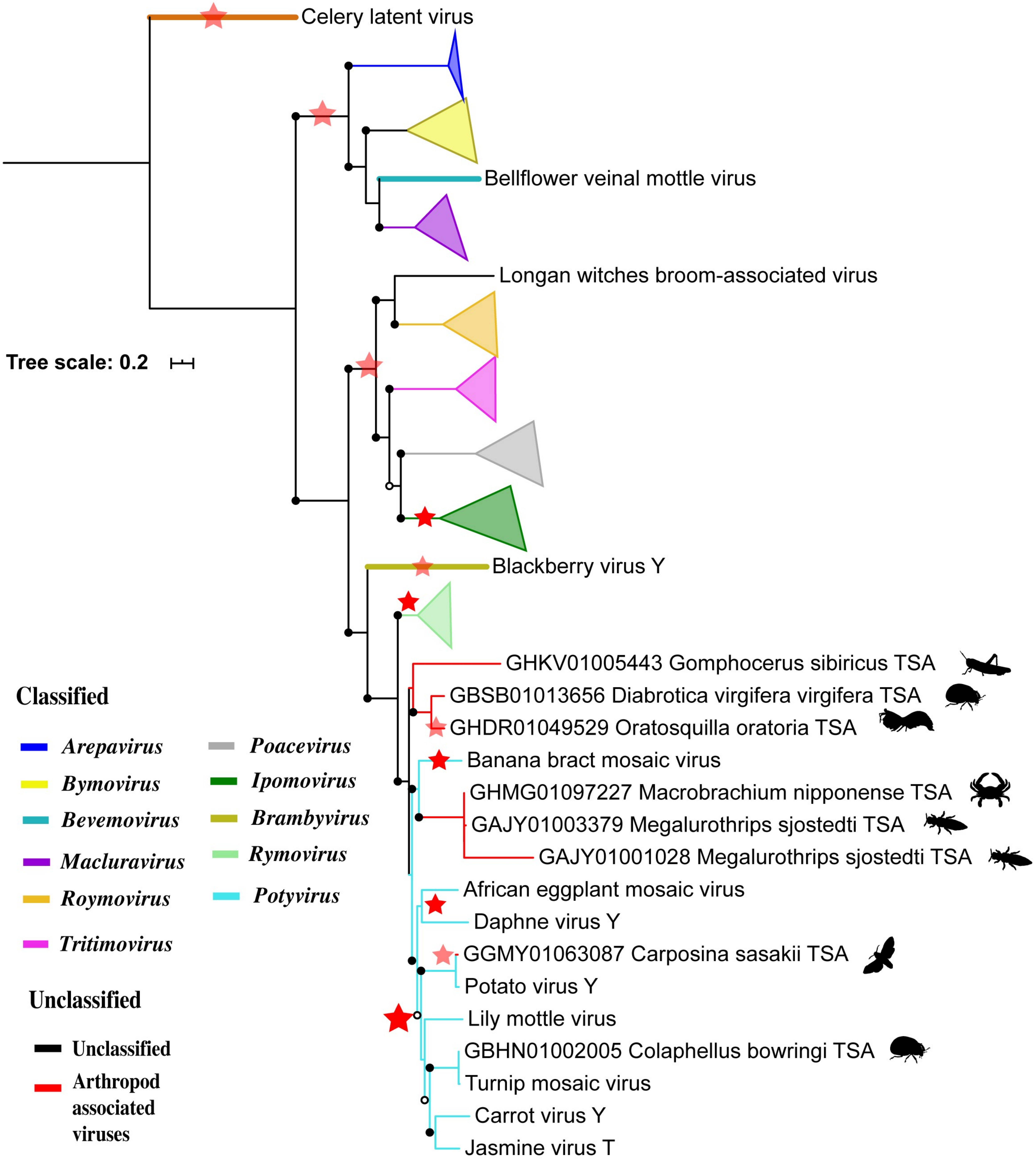


**Supplementary Figure 10: Phylogeny of viruses related to the *Potyviridae*..**

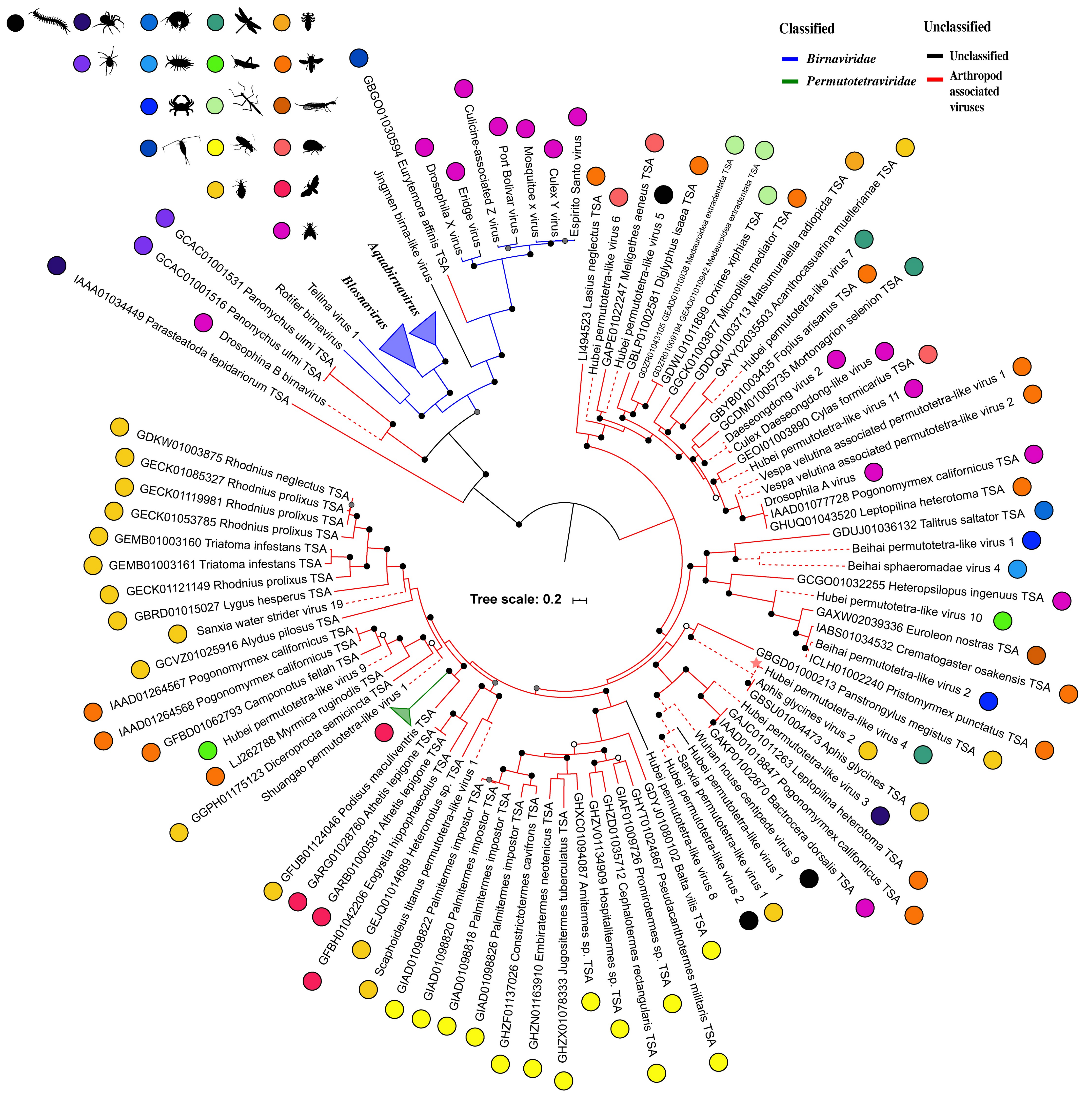


**Supplementary Figure 11: Phylogeny of viruses related to the "Birna-Permutotetra".**

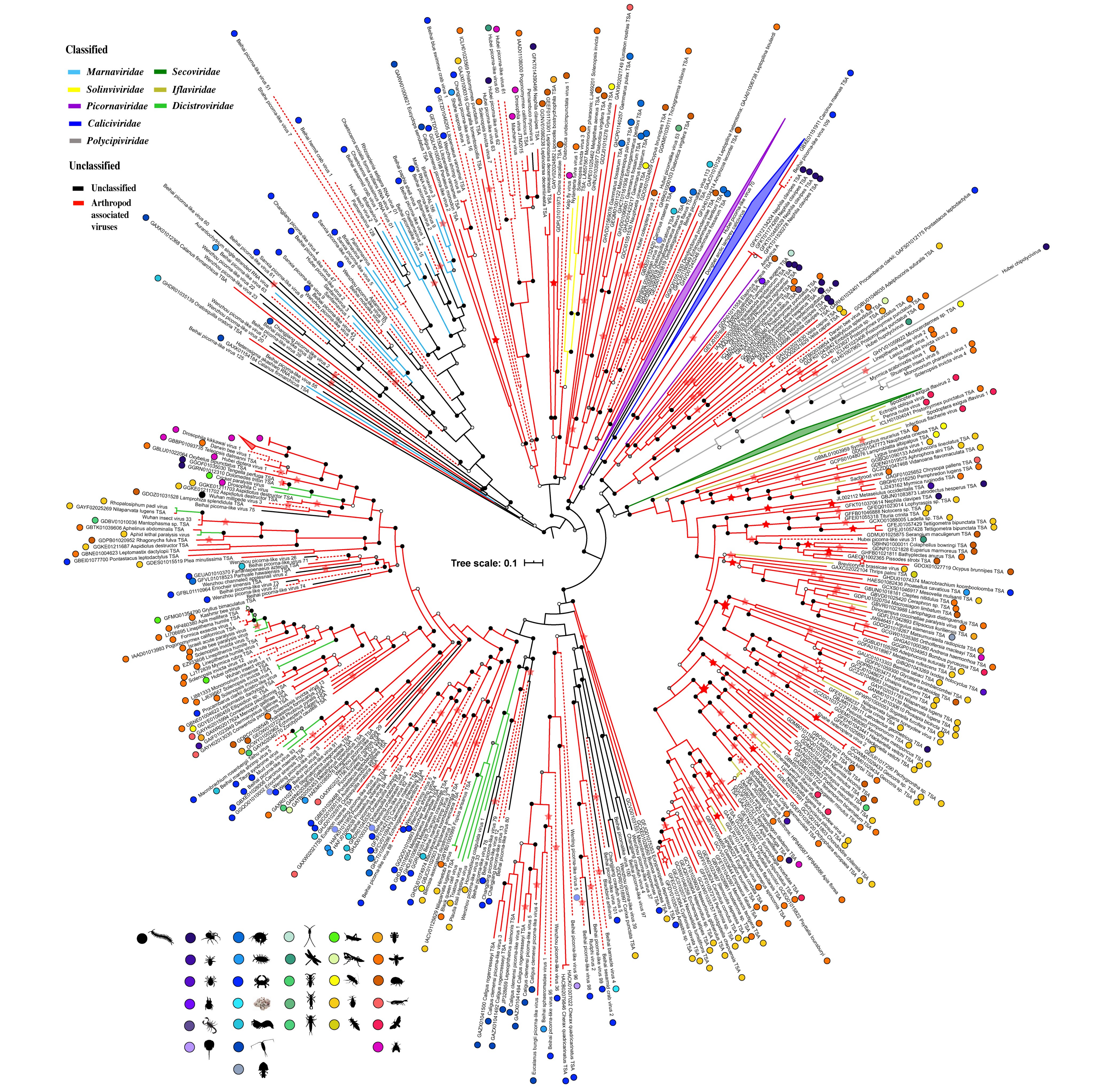


**Supplementary Figure 12: Phylogeny of viruses related to the *Picornavirales***

### ds RNA viruses

- S13: Phylogeny of viruses related to the “Ghabri-Botybri”.
- S14: Phylogeny of viruses related to the *Durnavirales*.
- S15: Phylogeny of viruses related to the *Reovirales*.
- S16: Phylogeny of viruses related to the *Partitiviridae*.

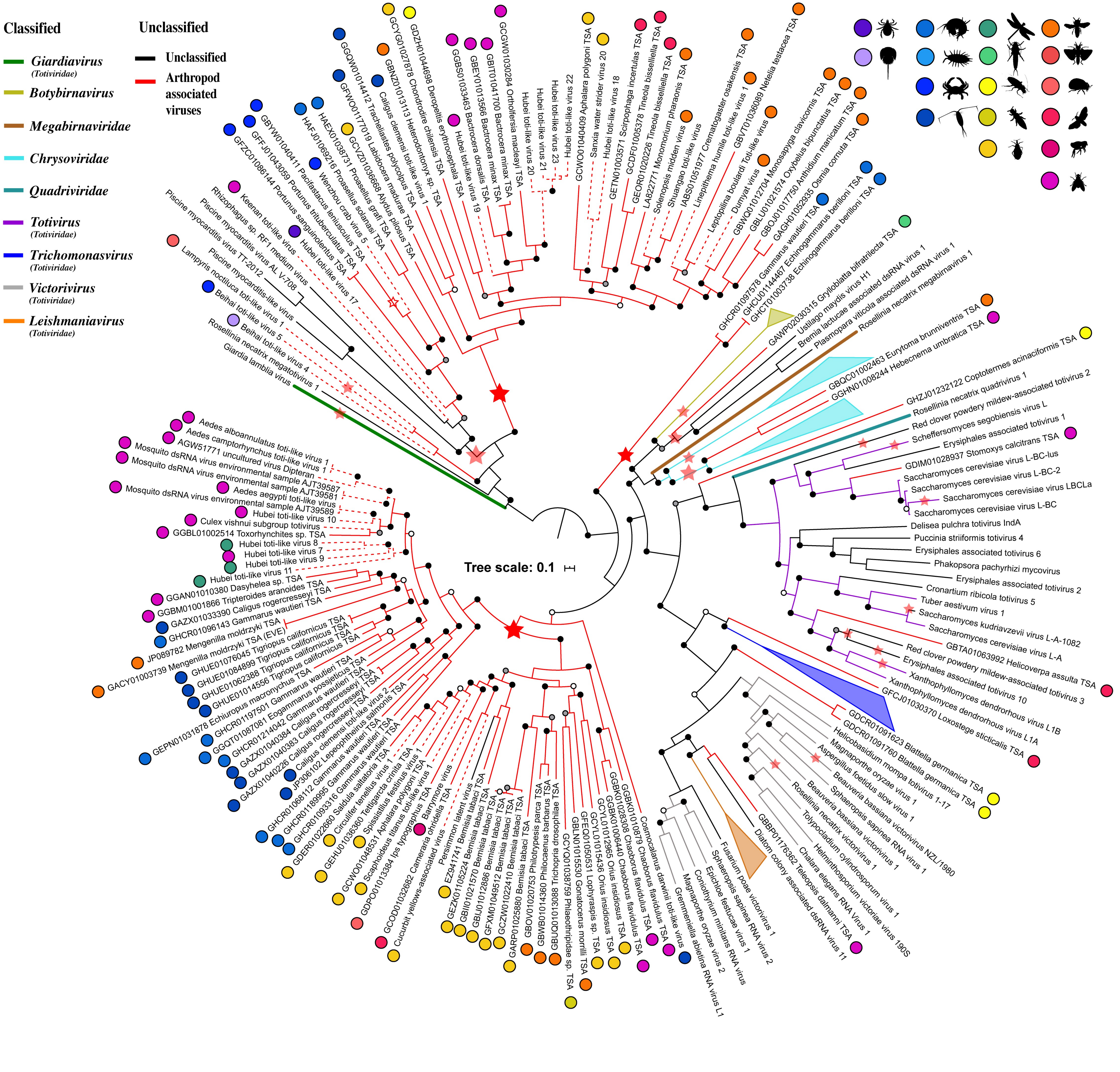


**Supplementary Figure 13: Phylogeny of viruses related to the "Ghabri-Botybri".**

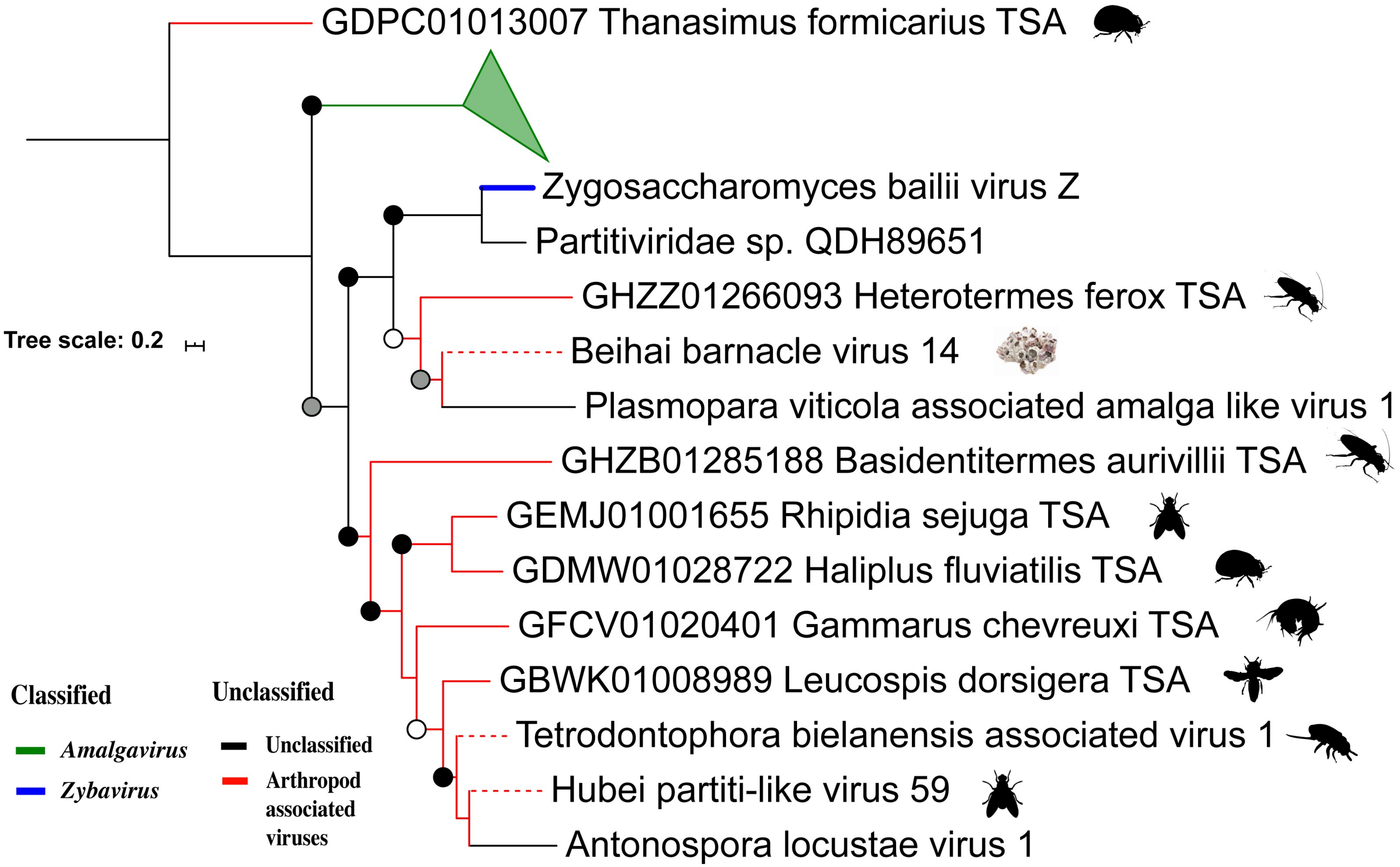


**Supplementary Figure 14: Phylogeny of viruses related to the *Durnavirales*.**

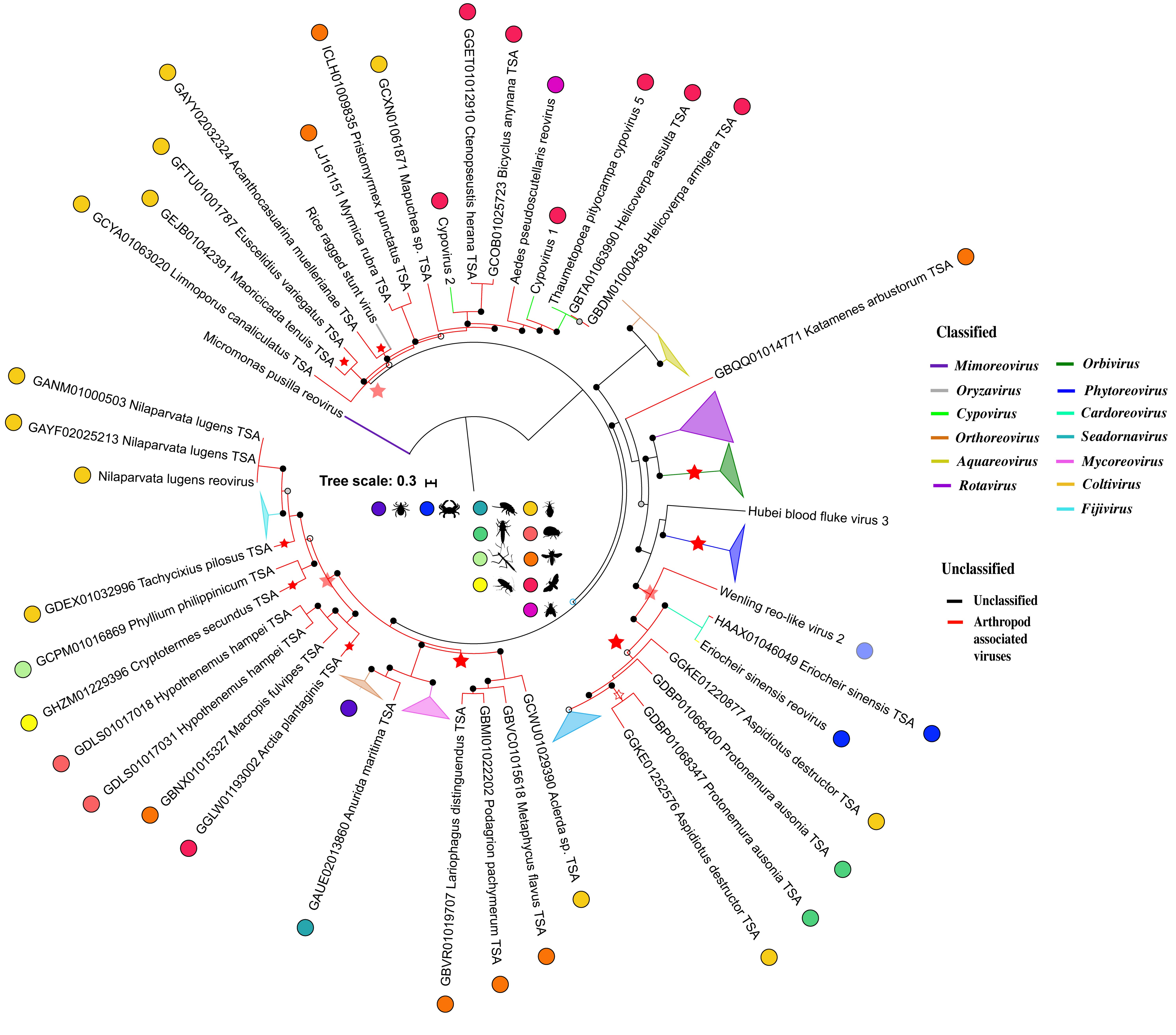


**Supplementary Figure 15: Phylogeny of viruses related to the *Reovirales*.**

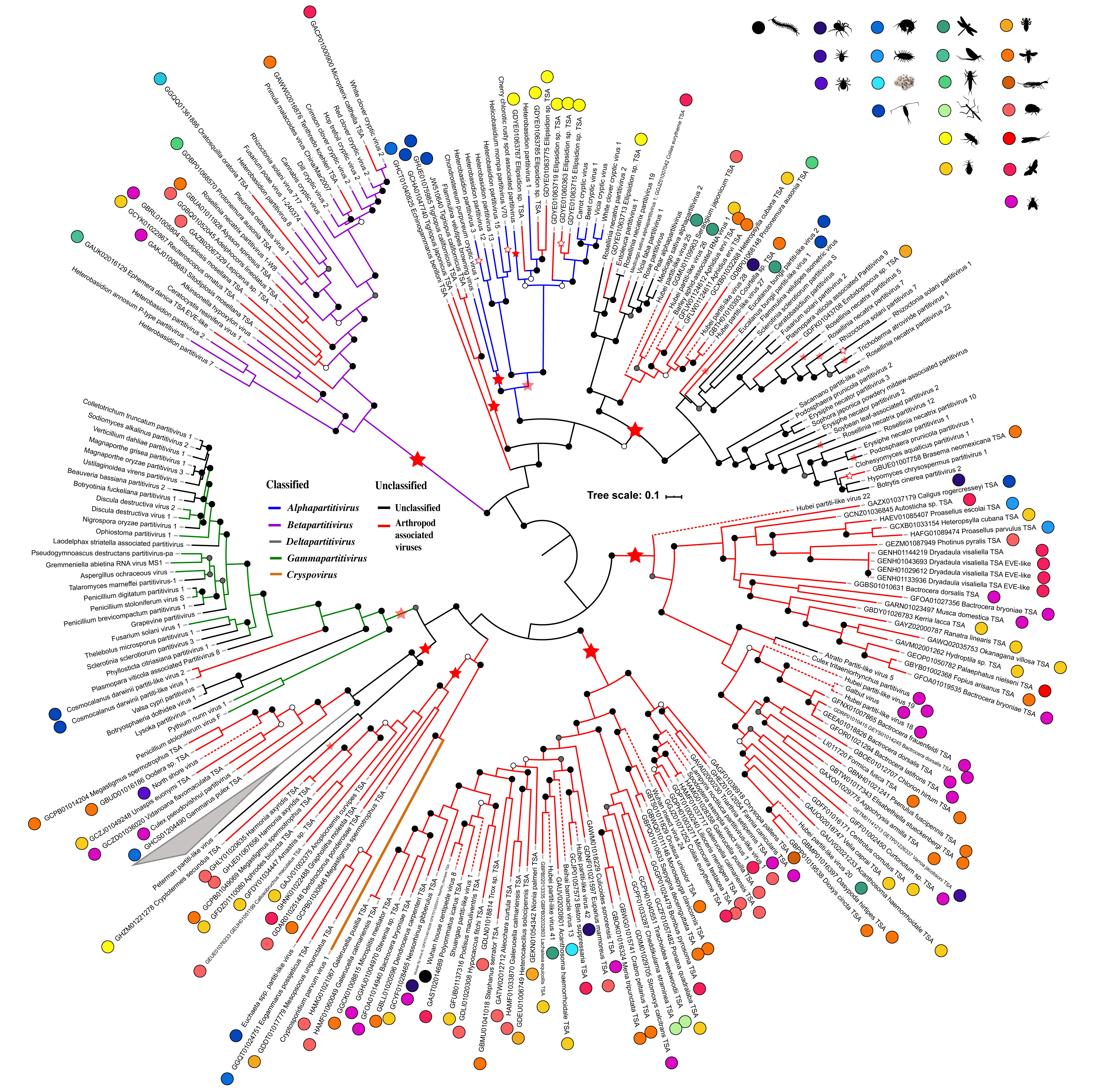


**Supplementary Figure 16: Phylogeny of viruses related to the *Partitiviridae*.**

### (-) ssRNA virus

- S17: Phylogeny of viruses related to the *Qinviridae*.
- S18: Phylogeny of viruses related to the *Chuviridae*.
- S19: Phylogeny of viruses related to the *Mononegvirales*.
- S20: Phylogeny of viruses related to the *Bunyavirales*.
- S21: Phylogeny of viruses related to the *Orthomyxoviridae*.

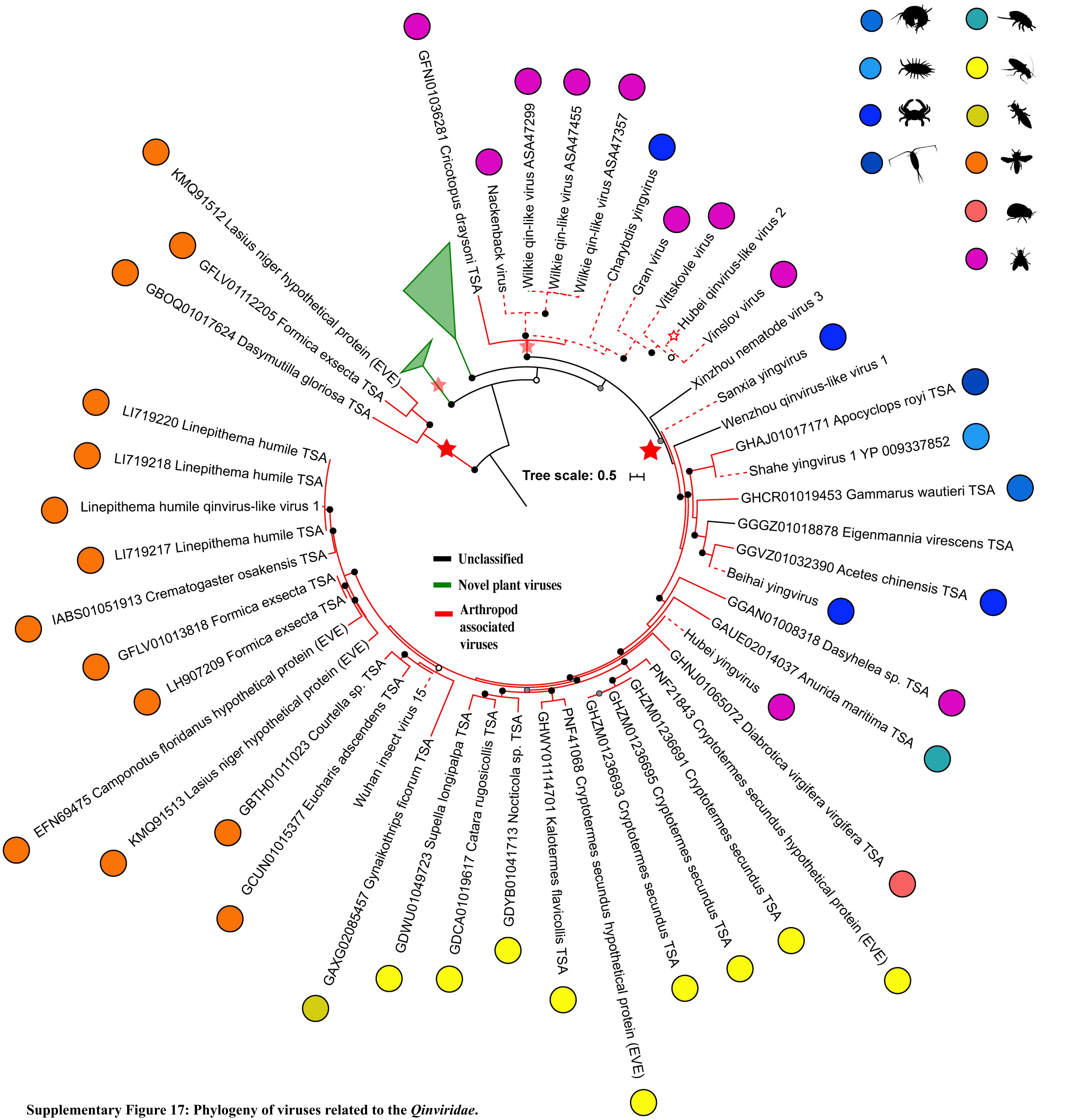


**Supplementary Figure 17: Phylogeny of viruses related to the *Qinviridae*.**

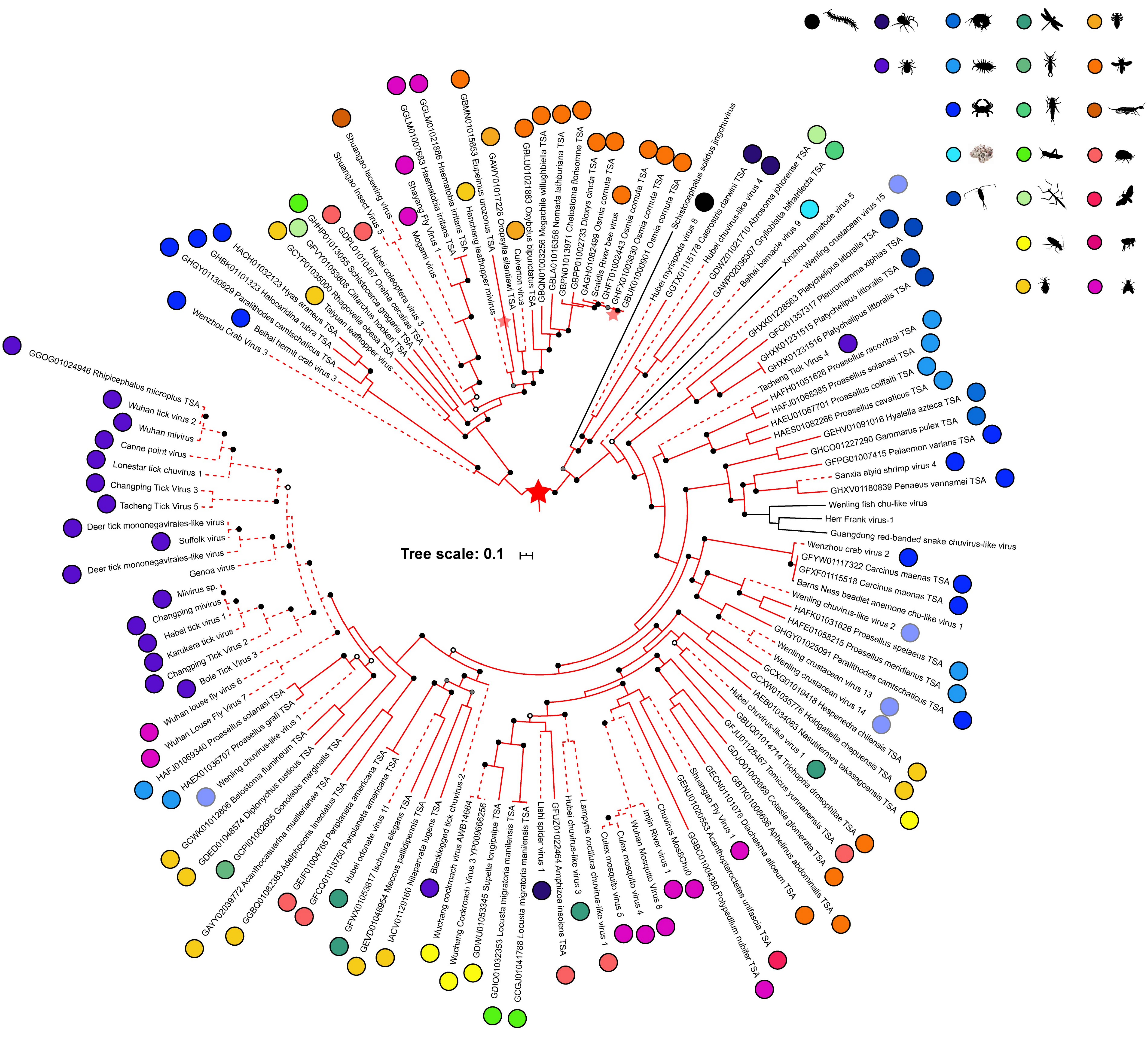


**Supplementary Figure 18: Phylogeny of viruses related to the *Chuviridae*.**

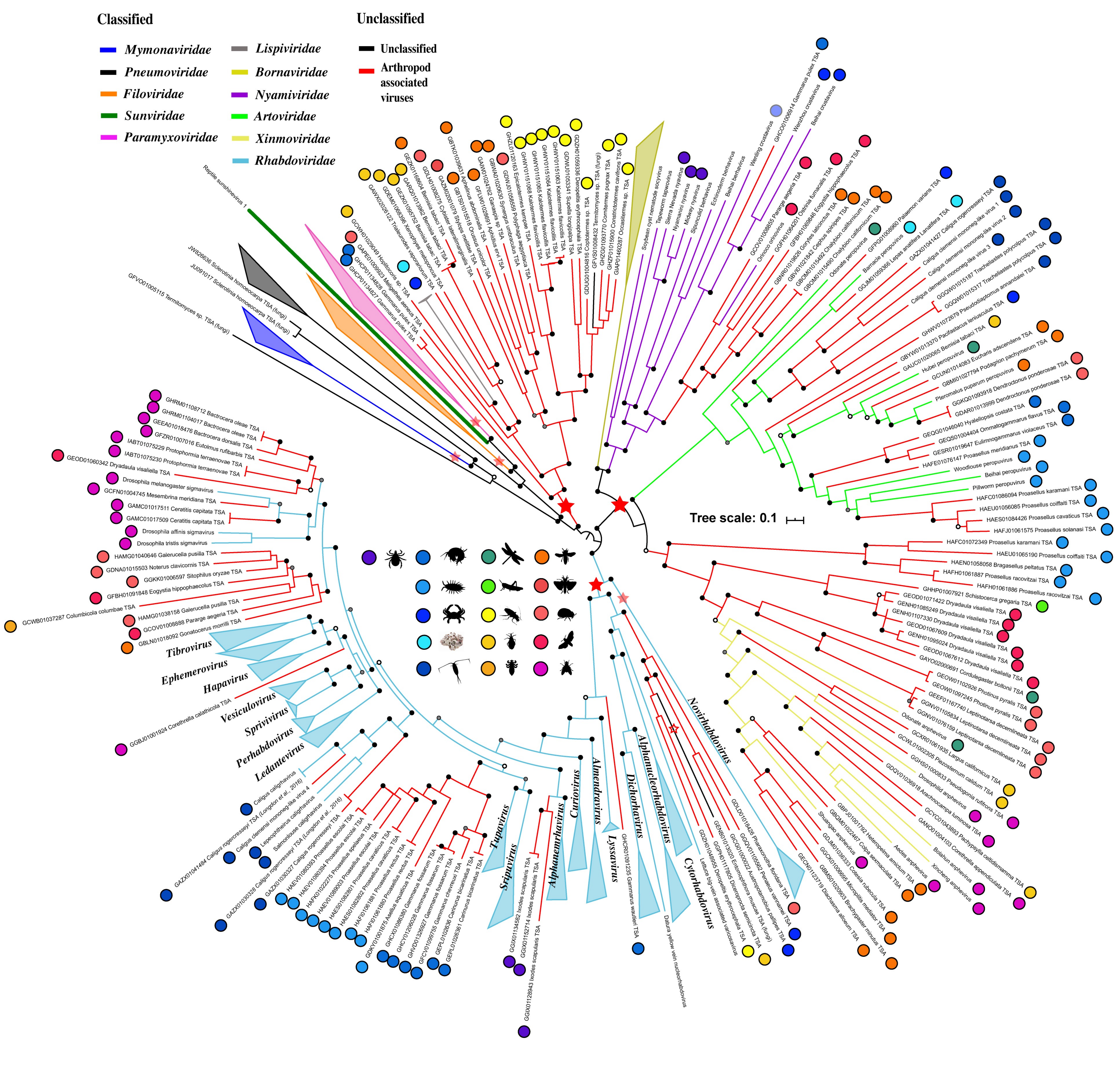


**Supplementary Figure 19: Phylogeny of viruses related to the *Mononegvirales*.**

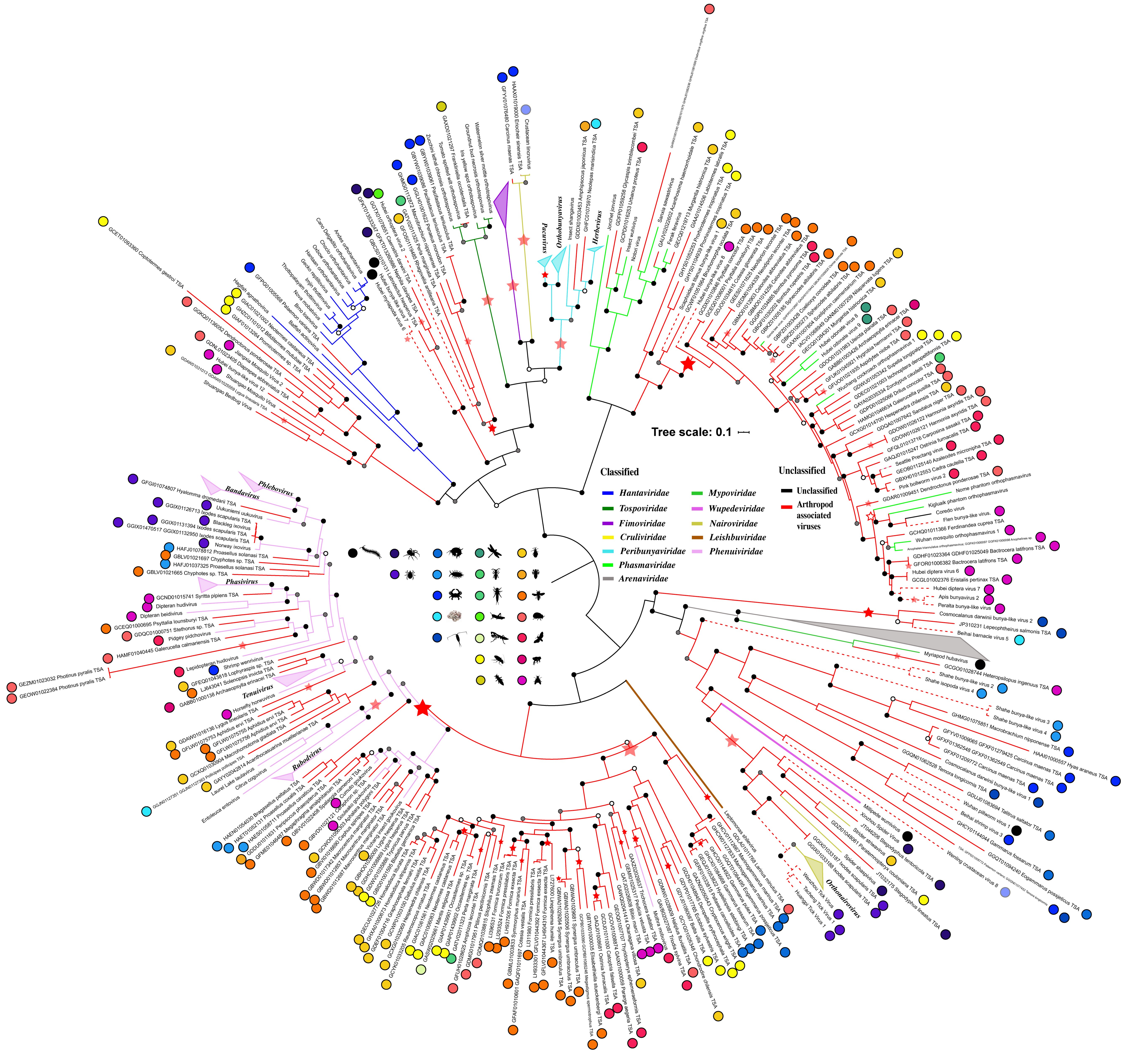


**Supplementary Figure 20: Phylogeny of viruses related to the *Bunyavirales*.**

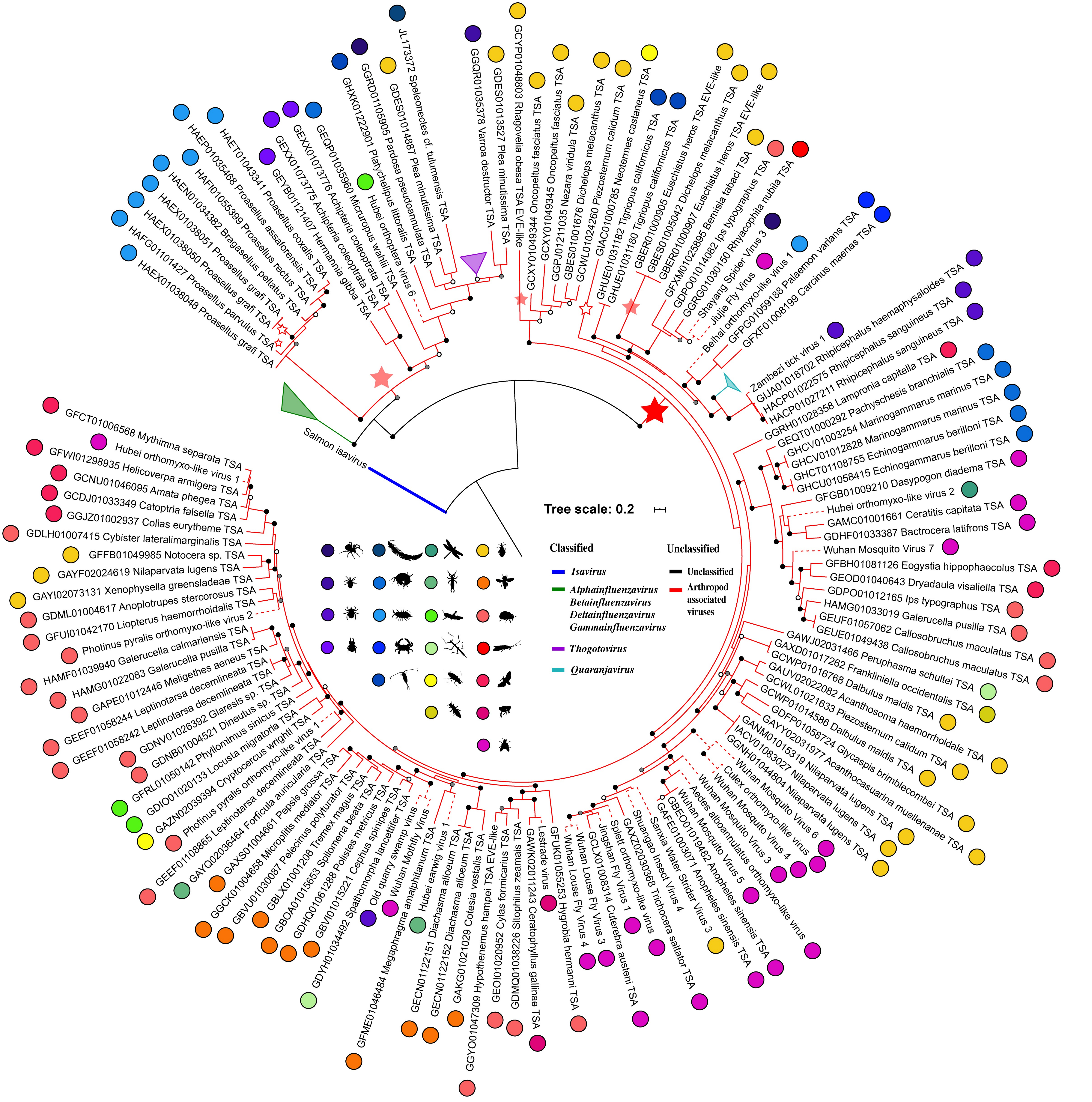


**Supplementary Figure 21: Phylogeny of viruses related to the *Orthomyxoviridae*.**
